# Proteomic signatures of cold adaptation and sponge host adaptation in marine Archaea

**DOI:** 10.64898/2026.11.02.763592

**Authors:** Kari Green, R. Eric Collins

## Abstract

Temperature shapes the biochemistry of microbial proteins, and several amino acid compositional indices have been proposed to detect cold adaptation in microbial proteomes. Whether these indices capture cold adaptation in Archaea, which are abundant in cold marine environments such as the Arctic Ocean, has not been tested. We calculated six amino acid compositional indices (arginine to lysine ratio, acidic residue proportion, aliphatic index, aromaticity, grand average of hydropathicity, and proline residue proportion) across global protein families for 142 Marine Group I (MG-I; Thaumarchaeota) and 146 Euryarchaeota genomes from the Bacterial and Viral Bioinformatics Resource Center (BV-BRC), together with three MG-I metagenome-assembled genomes (MAGs) recovered from the Western Arctic Ocean. The Arctic MAGs fell within the range of index and phylogenetic distances of the wider MG-I population, and no signature of cold adaptation was detected. Instead, index values differed between the two main MG-I phylogenetic groups and between freshwater and marine genomes, and the Euryarchaeota formed five index-based clusters consistent with phylogeny. A group of 19 MG-I MAGs associated with sea sponges had significantly higher arginine to lysine ratios than other MG-I, differed in the usage of all amino acids except alanine and lysine, and had distinct codon usage and higher GC content; their anomalous placement in the MG-I phylogeny is consistent with long-branch attraction. These results indicate that amino acid compositional indices developed for Bacteria have limited power to detect cold adaptation in Archaea, but reveal strong proteomic signatures associated with host adaptation in sponge-associated Thaumarchaeota.

## 1 Introduction

The TACK Superphylum is one of the most extensively studied clades today (Adam et al., 2017; Baker et al., 2020). Thaumarchaeota is thought to be one of the most numerically abundant clades of Archaea on the planet, nearing 20% of total microbes in some deep ocean and soil environments (Baker et al., 2020; Brochier-Armanet et al., 2008; Karner et al., 2001; Reji and Francis, 2020; Reji et al., 2019; Santoro et al., 2011; Spang et al., 2015). A highly studied representative of this group, *Nitrosopumilus maritimus*, has adapted to life under extreme nutrient limitations and high specific oxidation rates of ammonium at concentrations found in open oceans (Martens-Habbena et al., 2009). Currently, one phylum, the Euryarchaeota, does not fall into a superphylum but contains the greatest diversity of cultured lineages including members of the classes Methanopyri, Methanococci, Methanobacteria, Methanomicrobia, Archaeoglobi, Halobacteria, Thermococci, and Thermoplasmata (Baker et al., 2020; Offre et al., 2013). Euryarchaeota are broadly associated with diverse metabolisms including methanogenesis, methane-oxidation, denitrification, sulfate reduction, iron oxidation, and organotrophy (Offre et al., 2013). Culture-independent studies have shown that these organisms are important contributors to global biogeochemical cycling, such as the production of methane and the remineralization of nitrate (Baker et al., 2020; Offre et al., 2013). However, these classifications are always evolving. Rinke et al. (2020) proposed a major reclassification of the Domain consisting of 16 phyla, combining the TACK superphylum into one, and separating the Euryarchaeota into five new phyla. The first environmental archaeal sequences detected in marine environments were separated into two groups: Marine Group I (MG-I), now known as Thaumarchaeota, and Marine Group II (MG-II), within Euryarchaeota (Brochier-Armanet et al., 2008; Yilmaz et al., 2016). This structure has been further expanded to contain three main marine groups. Within Euryarchaeota, two groups were identified: Poseidoniales or MG-II (Pereira et al., 2019; Rinke et al., 2020), and Pontarchaea or Marine Group III (MG-III) (Baker et al., 2020; Yilmaz et al., 2016). Currently, MG-II representatives are not well understood as there are no cultured representatives from this group (Pereira et al., 2019). 16S rRNA gene sequences have revealed that MG-II may be divided into two main monophyletic groups, the MG-IIa, now known as *Candidatus* Poseidonaceae, and the MG-IIb, now known as *Candidatus* Thalassarchaeaceae (Rinke et al., 2020). The MG-III are commonly found in oceans worldwide; however, they are typically found in very low abundances. They have been shown to contain genes for recycling high molecular weight organic compounds (Baker et al., 2020), but the majority of their metabolic functions are unknown. For the remainder of the text, the archaeal clades of interest will be referred to by their marine group name (MG-I; MG-II; MG-III).

### 1.1 Ammonia oxidation by MG-I

Nitrogen is a necessary element for life and nitrogenous compounds are major limiting nutrients for marine primary production (Chang and Devol, 2009). Microbial nitrogen cycling is highly diverse but can be conceptualized as two main processes: nitrification (oxidation) and denitrification (reduction). Nitrification is a two-step process involving the oxidation of ammonia to nitrite, and nitrite to nitrate in aerobic environments (Figure 1) (Martens-Habbena et al., 2009). Aerobic ammonia oxidation is the first rate-limiting step in the nitrification process catalyzed by ammonia-oxidizing microbes such as MG-I Archaea (Martens-Habbena et al., 2009). Aerobic ammonia oxidation is the only oxidative biological process that links reduced and oxidized pools of inorganic nitrogen in nature (Martens-Habbena et al., 2009). The transformation of nitrogen directly influences primary producers who may differ in their ability to utilize different nitrogen compounds (Ward, 2005). It has been shown that MG-I possess ammonia monooxygenase genes (*amoA*) that oxidize ammonium to nitrite (Baker et al., 2020; Liu et al., 2018). These organisms have been shown to oxidize ammonia at low concentrations (Martens-Habbena et al., 2009) and obtain ammonia from urea and cyanate found in the open ocean (Baker et al., 2020; Christman et al., 2011; Liu et al., 2018; Palatinszky et al., 2015). Ammonia oxidation rates are often maximized near the bottom of the euphotic zone and are likely influenced by light levels in shallower waters where the activity of ammonia-oxidizing archaea (AOA) is minimized (Ward, 2005), suggesting light inhibition. There have also been suggestions that their distribution may be controlled by the availability of ammonia as their summer blooms coincide with high nitrite summer peaks (Horak et al., 2013; Liu et al., 2018; Martens-Habbena et al., 2009), as observed off the coast of Georgia, USA where MG-I 16S ribosomal RNA genes and archaeal *amoA* genes coincided with peak nitrite concentrations (Liu et al., 2018). In every environment, there are two possibilities for ammonia consumption: nitrification or assimilation. Ammonia assimilation has been shown to favour increases in temperature compared to nitrification suggesting that as the Arctic continues to warm, assimilation may become more dominant (Baer et al., 2014; Ward, 2005). Nitrification rates and total *amoA* gene abundance have been found to be much higher in winter months than in summer which was attributed to the cold and dark conditions which increase the chemoautotrophic potential of the community (Christman et al., 2011). Baer et al. (2014) demonstrated that nitrification rates were up to 2-fold greater than ammonia assimilation in the ice-covered winter and spring seasons and appear to be driven by a high concentration of ammonia in the environment. Although ammonia oxidation and assimilation were not found to be significantly correlated, there has been an inverse relationship noted where the highest nitrification rates observed were at depths where the lowest assimilation rates were occurring (Ward, 2005). Phytoplankton have been shown to outcompete bacterial and archaeal nitrifiers for ammonia when waters are warmer and light penetration is higher (Baer et al., 2014; Martens-Habbena et al., 2009; Ward, 2005). During the winter, this competition is reduced as primary production is limited by light and bioavailable nutrients under ice coverage (Baer et al., 2014; Oziel et al., 2022). Nutrient limitation is speculated to become a more prominent issue with decreased sea-ice coverage allowing more light to penetrate the water column (Oziel et al., 2022). This is likely to be one of the main controls on ammonia oxidation productivity. However, among bacteria, true psychrophiles (organisms that grow exclusively at low temperatures) have been shown to be less sensitive to substrate concentrations and have the potential to remain competitive against other phytoplankton for ammonia (Baer et al., 2014; Yager and Deming, 1999). Environmental factors such as light, competition, substrate, and community composition all support the process of nitrification and the use of ammonia as an energy source in the winter and spring months (Baer et al., 2014). As the Arctic continues to warm, it is expected that light penetration will increase, primary production rates will increase, begin earlier, and end later, and thus it is likely that ammonia competition will become greater. However, if the nitrifiers residing in these environments are “true-psychrophiles”, they may be able to maintain the same levels of energy production even when concentrations are reduced.

**Figure 1:**
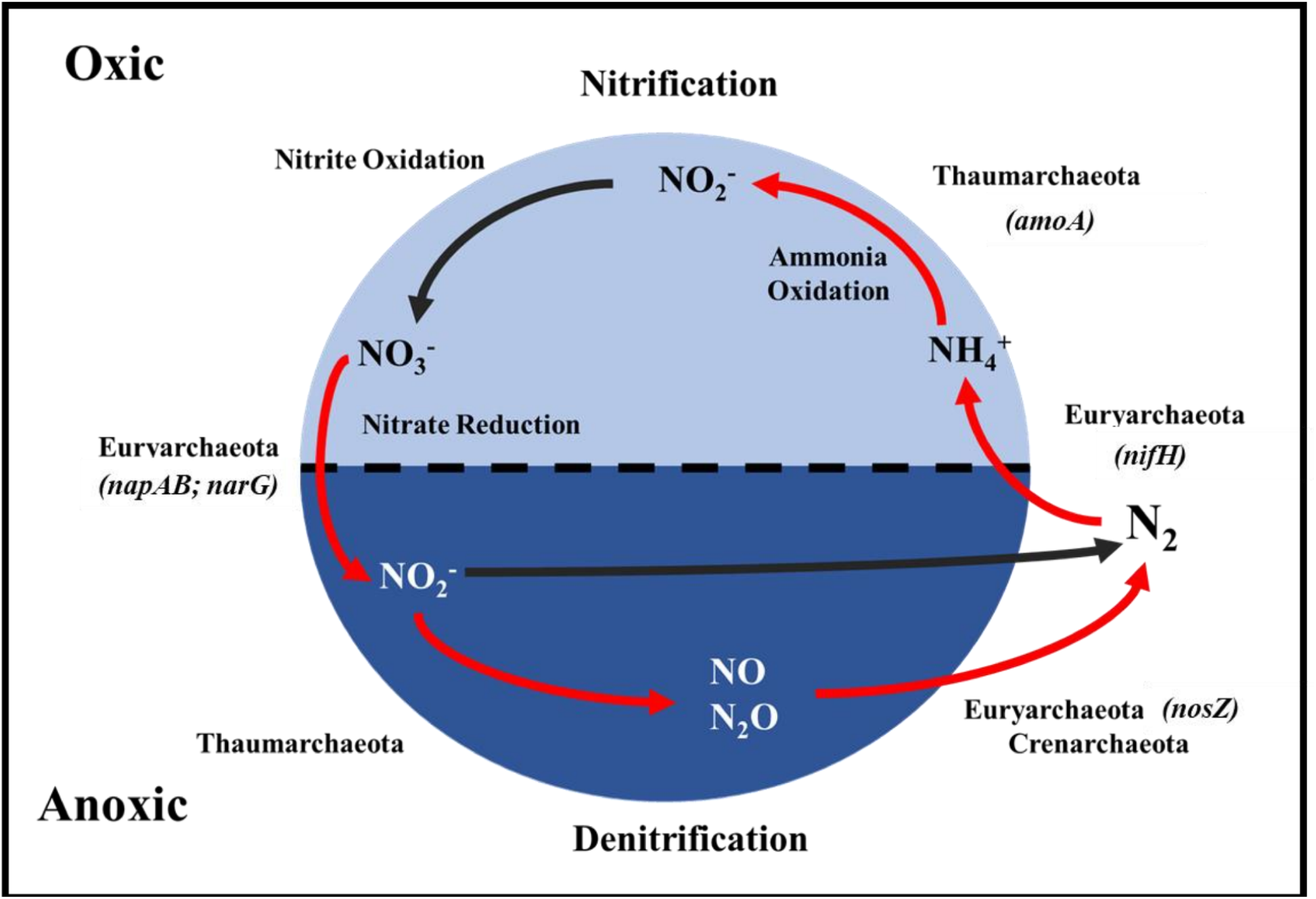
The process of nitrification and denitrification occurring in the ocean. Portions of the cycle in light blue occur in an oxic environment, while dark blue indicates an anoxic environment. Red arrows signify steps within the process that Archaea are believed to play a role in and if known, the main archaeal gene involved is listed. The genes are as follows: *amoA*: ammonia monooxygenase (PGF_02515073); *nifH* : nitrogenase iron protein (PGF_00025954); *napA*: periplasmic nitrate reductase (PGF_00031036); *napB*: nitrate reductase electron transport subunit (PGF_00025698); *narG*: nitrate reductase (PGF_09372748); *nosZ* : nitrous oxide reductase (PGF_00026089) (Yilmaz et al., 2016).

### 1.2 Archaeal cold adaptation

Temperature greatly impacts cellular biochemistry because many chemical reactions are limited by diffusion of the reagents to the active site of the enzyme that catalyzes the reaction. In warmer environments chemical processes are sped up, but if the temperature is too high, proteins denature, and cells die. In colder environments, processes slow down, and protein flexibility is greatly hindered. All microorganisms are at a temperature equilibrium with their environment, therefore organisms living in extreme temperatures, like those seen in the Arctic Ocean, typically have genetic and proteomic adaptations to withstand the conditions. Psychrophilic microorganisms can grow at −10 to 15 °C, metabolize at −20 °C, and survive at −45 °C (Cavicchioli, 2006). Psychrophiles typically have decreased enzymatic reaction rates and increased osmotic and oxidative stress (Cimen et al., 2020; Goordial et al., 2015). These organisms tend to have protein adaptations that enable them to live in sub-zero environments which allow DNA replication, transcription, translation, and membrane fluidity to be maintained (Goordial et al., 2015). Some adaptations in protein structure can include fewer salt bridges, proline residues, and hydrogen bonds at low temperatures (Goordial et al., 2015). In general, cold-active enzymes will have fewer strong amino acid residue bonds (Cimen et al., 2020). They also tend to increase the copy number of core genes to combat the low reaction rates brought on by colder temperatures (Goordial et al., 2015). The model cold-adapted archaeon *Methanococcoides burtonii* has many molecular adaptations associated with psychrophilia including differences in its Elongation Factor 2 (EF2) (Cavicchioli et al., 2002; Thomas and Cavicchioli, 2000), changes in its intracellular levels of potassium glutamate and potassium aspartate (Cavicchioli et al., 2002), and a higher rate of membrane-lipid unsaturation (Nichols et al., 2004). Generally, proteins from cold adapted Archaea have more non-charged amino acids (Gln and Thr), and fewer hydrophobic residues (Leu) (Cavicchioli, 2006; Saunders et al., 2003). These changes are compensations that increase flexibility, reduce the activation energy of the protein-substrate transition state, and increase catalytic efficiency at low temperatures (Cavicchioli, 2006; Saunders et al., 2003). Phylogenetically, organisms from cold environments are typically more closely related to each other than to their thermophilic counterparts. For example, MG-I members such as *N. maritimus* identified in low-temperature environments were more closely related to each other than to thermophilic members of the same clade (Cavicchioli, 2006). Overall, the responses to cold environments in Archaea and Bacteria are quite similar (Goordial et al., 2015). It is not currently feasible to experimentally determine the optimal growth temperature (OGT) of every species in Arctic seawater, but there are several metrics that can be used to infer OGT by identifying differences in genomic, transcriptomic, and proteomic composition using -omics techniques. Several indices have been suggested as a measure of cold adaptation in proteins including arginine to lysine ratio, proline residue abundance, grand average of hydropathicity (GRAVY), aromaticity, aliphatic index, and acidic residue abundance (Goordial et al., 2015). Each index is assigned a score that can be used to determine whether the protein is cold adapted. It is expected that these scores will be lower in cold adapted organisms. A lower arginine to lysine score indicates a preference for using lysine, which is smaller and requires fewer nitrogen atoms. GRAVY is the measure of hydropathicity of each amino acid. It is expected that amino acids that are less hydrophobic and amino acids with smaller aliphatic side chains would allow for increased flexibility. The environmental conditions preferred by individuals or populations of any microbial taxon are encoded in its genome, yet predicting those preferences from genomic data is an unsolved problem for many environmentally relevant parameters, including temperature and salinity. For example, predictions of optimal growth temperature in Bacteria and Archaea using only genomic information are robust for temperatures above 37 °C, but poor at lower temperatures (Li et al., 2019). The benefits of being able to make such predictions would be widespread, including incorporating rates of microbial growth and metabolism into numerical models of ocean biogeochemistry based on easily accessible metagenomic sequencing. A number of genomic targets have been considered for such predictions in the literature, including nucleotide base composition, amino acid usage, and gene content. To assess cold adaptation in Archaea, a collection of archaeal metagenome-assembled genomes (MAGs) and genomes from cultured isolates were obtained from the BV-BRC database. Additional MAGs assembled from Arctic Ocean metagenomes in a companion study (Green and Collins, 2026) were included in the collection. Metadata and sequences including nucleotide, amino acid, and coding regions were downloaded. The genomes obtained from the BV-BRC portal were of diverse origin encompassing fresh water (lakes, rivers, ponds, and river sediment), marine water (oceans, seas, marine sediment, and saltwater aquariums), or other (wastewater, soil, feces, and brackish water). These samples were also collected from various years, seasons, and locations. Using the amino acid sequences of the annotated genes, six amino acid indices were calculated: arginine to lysine ratio, aliphatic index, aromaticity, acidic residue proportion, GRAVY index, and proline residue proportion.

### 1.3 MG-I host adaptation in sea sponges

Sponges are one of the first groups of organisms that evolved to become multicellular (Kenny et al., 2020). Most have a highly conserved body structure which consists of channels that filter water, up to thousands of liters daily, using pumping cells (Kenny et al., 2020; Slaby et al., 2017). Sponges have very large genomes, approximately double that of humans (Kenny et al., 2020). The sponge microbiome is known to include as many as 52 different microbial phyla, however the type and amounts may vary between species (Slaby et al., 2017; Webster and Thomas, 2016). There can be thousands of symbiont lineages associated with a single sponge, typically dominated by Proteobacteria (mostly Alpha- and Gammaproteobacteria), Acidobacteria, Actinobacteria, Chloroflexi, Cyanobacteria, Crenarchaeota, as well as symbionts of several candidate phyla (Webster and Thomas, 2016). The sponge-associated microbes can perform carbon, nitrogen, and possibly sulfur and vitamin metabolism (Bayer et al., 2008; Hentschel et al., 2012; Horn et al., 2016). The microbiome of sponges has been shown to preferentially express genes related to cellular defense (restriction-modification systems, toxin-antitoxin systems, host colonization, and extracellular matrix utilization) (Slaby et al., 2017). This observation is likely due to the extensive water filtering performed by sponges, which increases the contact with potential pathogens, toxins, and other stress factors (De Goeij et al., 2009; Slaby et al., 2017). Generally, it has been observed that sponge symbiont genomes are larger than those of free-living organisms and contain a higher GC-content (50-60%) (Horn et al., 2016; Slaby et al., 2017). Enrichment of certain systems, such as amino acid transport and metabolism, replication, recombination, and repair were seen in the symbionts, while transcription, signal transduction mechanisms, cell wall and membrane biosynthesis, and cell mobility were depleted (Slaby et al., 2017). These findings were all found to occur in Bacteria, while no archaeal analysis was performed. A large issue with identifying these traits in Archaea is the lack of high-quality annotations available for archaeal proteins. The genetic changes seen in sponge symbionts are likely due to increased exposure to free DNA and other pathogens the microbes encounter during sponge filtration and reflect an evolutionary pressure toward host adaptation. High levels of horizontal gene transfer (HGT) are likely occurring and high GC-content and defense mechanisms appear to be a requirement for sponge adaptation (Horn et al., 2016).

### 1.4 Objectives

The objective of this study is to identify signatures of cold adaptation in the amino acid usage and gene content of Archaea, to work towards future genomic predictions of optimal growth temperature in these important marine microbes. Based on previous work in cold adapted Bacteria (Gentilhomme, 2020; Goordial et al., 2015; Raymond-Bouchard et al., 2018), we hypothesized that cold adaptation would be apparent in the genomes of cultured and uncultured Archaea at the proteomic level when measured with six amino acid usage indices, and that environmental temperature would be a factor influencing the amino acid usage in Archaea, and finally that phylogeny would reflect the environmental niches of Archaea.

## 2 Materials and Methods

### 2.1 Genome dataset

The dataset comprised 142 MG-I (Thaumarchaeota) genomes and MAGs and 146 Euryarchaeota genomes and MAGs obtained from the BV-BRC portal (Figures 2 and 7), originating from diverse environmental sources, collection years, seasons, and locations. Collection sources were categorized as fresh water (lakes, rivers, ponds, and river sediment), marine water (oceans, seas, marine sediment, and saltwater aquariums), or other (wastewater, soil, feces, and brackish water). Outgroup genomes were included in each analysis (Sections 3.1 and 3.2). Three MG-I MAGs recovered from the Western Arctic Ocean during the 2015 GEOTRACES expedition (bins 29, maxbin.015_sub, and maxbin.074_sub; Green and Collins, 2026) were added to the MG-I dataset. Completeness and contamination of MG-I genomes were estimated with CheckM (Parks et al., 2015) where these values were not available in the genome metadata (Supplementary Table S1).

**Figure 2:**
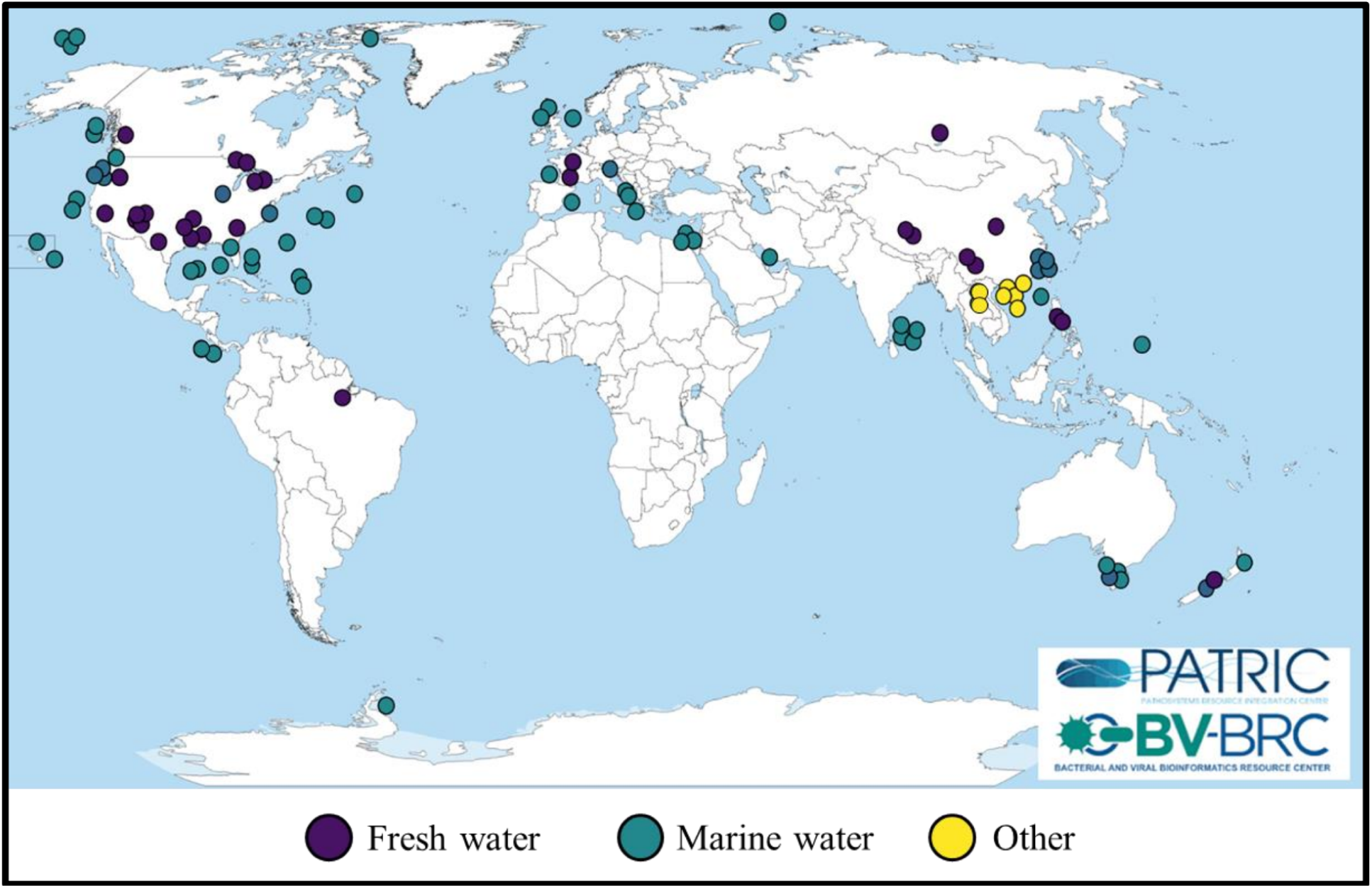
Geographical map showing the sampling locations of the MG-I genomes obtained from the BV-BRC database. Colours of each circle indicate the sampling source: fresh water (purple), marine water (teal), and other (yellow).

### 2.2 Calculation of amino acid compositional indices

Archaeal genomes and metadata were collected and downloaded from the BV-BRC portal (https://www.bv-brc.org). The genomes were all classified as “good” in the database. Cold adaptation indices were calculated using existing scripts developed at the University of Alaska Fairbanks (https://github.com/agentilhomme/Genomic-Signatures-of-OGT-2020-). Briefly, global and local protein family assignments for each gene in each genome were downloaded. Using the amino acid sequences of each protein, statistical analysis of cold adaptation as a function of phylogeny was done in R (R Core Team, 2021) using RStudio. For each protein in each genome, the amino acid indices were calculated. The proteins were averaged based on their global protein family assignments. The indices were then standardized using Z-scores, which are numerical measurements that describe the relationship of the value to the mean and are measured in terms of standard deviations from the mean. The analysis was based on the phyla of Archaea and the Ward distances of the global protein families. Ward’s distance method uses a minimum variance criterion where the total within-cluster variance is minimized. At each step, this method finds the pair of clusters that results in the smallest increase in total within-cluster variance when merged. Statistical analysis was performed comparing various groups (phylogenetic, environmental, etc.) for each index using *t*-tests as all indices followed a normal distribution and had unknown variances. The null hypothesis assumed that the two groups in comparison have equal variances. The null hypothesis was rejected when the *P* -value *<* 0.05 indicating that the data readings were strong and probably not due to chance. The type of *t*-test used was an equal-variance independent *t*-test because the number of samples (genomes) in each group being compared was different. Degrees of freedom were calculated as *n*_1_ + *n*_2_ − 2.

### 2.3 Calculation of codon and amino acid usage

Sequences of all the genomic features (protein sequences and open reading frame DNA sequences) for each genome used in amino acid compositional indices analysis were downloaded from the BV-BRC database. Calculations of total amino acid usage by each genome were totaled in R. Codon usage analysis was performed in R using the seqinr package (Charif and Lobry, 2007). Briefly, codons within each open reading frame DNA sequence were counted, where a codon represents three DNA bases that correspond to an amino acid during protein translation. Total codon usage was presented either as a tally of each codon used in that genome or as a proportion of all the codons identified in that genome. Statistical analysis was performed comparing the sponge-associated genomes against the general population for the overall amino acid usage using *t*-tests. The null hypothesis assumed that the two groups in comparison have equal variances. The null hypothesis was rejected when the *P* -value *<* 0.05 indicating that the data readings were strong and probably not due to chance. The type of *t*-test used was an equal-variance independent *t*-test because the number of samples (genomes) in each group being compared was different. Degrees of freedom were calculated as *n*_1_ + *n*_2_ − 2.

## 3 Results

### 3.1 Amino acid compositional indices show little variation among MG-I MAGs

A collection of 142 MG-I MAGs was obtained from the BV-BRC database to assess the six established amino acid indices. The source and location of the MAGs can be seen in Figure 2. The genomes were of diverse sources including season, year, environment, and location. The majority of these samples were obtained from mid-latitude regions, with few from the poles. This is due to the lack of available archaeal data in polar regions, exemplifying the importance of recovering archaeal MAGs from polar regions such as the Western Arctic Ocean (Green and Collins, 2026). The individual indices for each protein were averaged across global protein families within each MAG (Figure 3). The outgroups (*Thermococcus alcaliphilus* AEDII12; *Halorientalis regularis* TNN28; *Candidatus* Bathyarchaeota strain NP77; *Candidatus* Bathyarchaeota strain B24-2; *Nanoarchaeum equitans* Kin4-M; Nanoarchaeota strain JGI OUT-1) showed distinct differences in indices, specifically in the arginine to lysine ratio and acidic residue proportion compared to MG-I. Little variation was observed across the MG-I for each index when standardized (Figure 3). The index distributions for the three MG-I genera used in this analysis can be seen in Supplementary Figure S1. In addition, the environmental niches which they occupied were not consistent with the phylogeny. Archaeal MG-I MAGs collected from the Western Arctic Ocean (Green and Collins, 2026) were also added to this analysis and included bins 29, maxbin.015_sub, and maxbin.074_sub. These MAGs were chosen as their completeness was above 70%; although lower than the MIMAG standard of 90%, they were the highest quality bins from the identified MG-I MAGs and were still considered to be medium-quality drafts. To investigate any differences between the Arctic MG-I MAGs and the general dataset, the index distances and the phylogenetic distances of each MAG compared to another MAG in the dataset were plotted in Figure 4. The indices for these MAGs were all within the normal distributions of the indices for the population tested. The bins did have slightly elevated distances in both the indices and phylogenetics, but these differences were not significant. Although there were not any detectable signatures of cold adaptation when comparing the Arctic MAGs to the general archaeal population, there were differences observed in the indices tested based on two other factors: phylogeny and environment. There were two main clades of MG-I (Supplementary Figure S2) encompassing the genera *Nitrosopumilus* and *Nitrosomarinus* (Group A) and *Nitrosarchaeum* (Group B) which were identified based on phylogenetic tree structure. It is important to note that there was a collection of *Nitrosopumilus* MAGs associated with sea sponges that clustered phylogenetically with Group B. The phylogenetic tree building tool used single copy genes to measure the relatedness between the different MAGs. Many of the MAGs included in this analysis did not have completeness or contamination measurements included in their metadata. CheckM was used to assess bin quality and can be seen in Supplementary Table S1. This is discussed further in Section 3.3. Between these two phylogenetic groups (*n*_1_ = 65 (Group A); *n*_2_ = 76 (Group B); degrees of freedom = 139), it was observed that there was a significant difference in the indices for the arginine to lysine ratio (*P <* 0.0001), acidic residue (*P <* 0.0001), GRAVY (*P <* 0.0001), and proline residue proportion (*P <* 0.0001) indices (Figure 5). Group B was found to have higher scores for all of the significantly different indices, excluding the acidic residue proportion. These differences were determined to be due to the outlier Nitrosopumilaceae MAGs in Group B identified above. In addition, there was also a collection of MAGs (2202732.58, 2202732.18, 2202732.35, 2202732.59, 2202732.61, and 2202732.62) that were not associated with sponges that were also members of this outlier Nitrosopumilaceae group, although they were not identified at the genus level. Their completeness and contamination scores ranged from 77.02–95.15% and 0–0.97%, respectively, suggesting that this observation was not caused by poor bin quality. With the removal of the outlier Nitrosopumilaceae MAGs, there was a lack of phylogenetic signal influencing the indices, which could speak to the evolutionary conservation of this group, which is further explored in Section 4.2. The second factor influencing variance within the indices was the collection environment. The majority of the MAGs (53%) were collected from marine environments, and it was observed that there were significant differences between the fresh and marine water MAGs in all of the amino acid indices tested in at least one of the phylogenetic groups (Group A: *n*_1_ = 9 (fresh water), *n*_2_ = 42 (marine water), degrees of freedom = 49; Group B: *n*_1_ = 28 (fresh water), *n*_2_ = 33 (marine water), degrees of freedom = 59). This observation was consistent within the two phylogenetic groups as the aliphatic (*P <* 0.0001), acidic residue proportion (*P <* 0.0001), GRAVY (*P <* 0.0001), and proline residue proportion (*P <* 0.0001) indices were significantly different in Group A, and the arginine to lysine ratio (*P <* 0.0001), aliphatic (*P <* 0.0001), aromaticity (*P <* 0.0001), and proline residue proportion (*P <* 0.0001) indices were significantly different within Group B when comparing the fresh water and marine water MAGs (Figure 6). The acidic residue proportion (significant in Group A), the arginine to lysine ratio (significant in Group B), and the proline residue proportion (significant in both groups) were lower in the MAGs from freshwater environments. The aliphatic and GRAVY indices were lower in the MAGs from marine environments. The aromaticity index was also lower in the marine MAGs in Group B.

**Figure 3:**
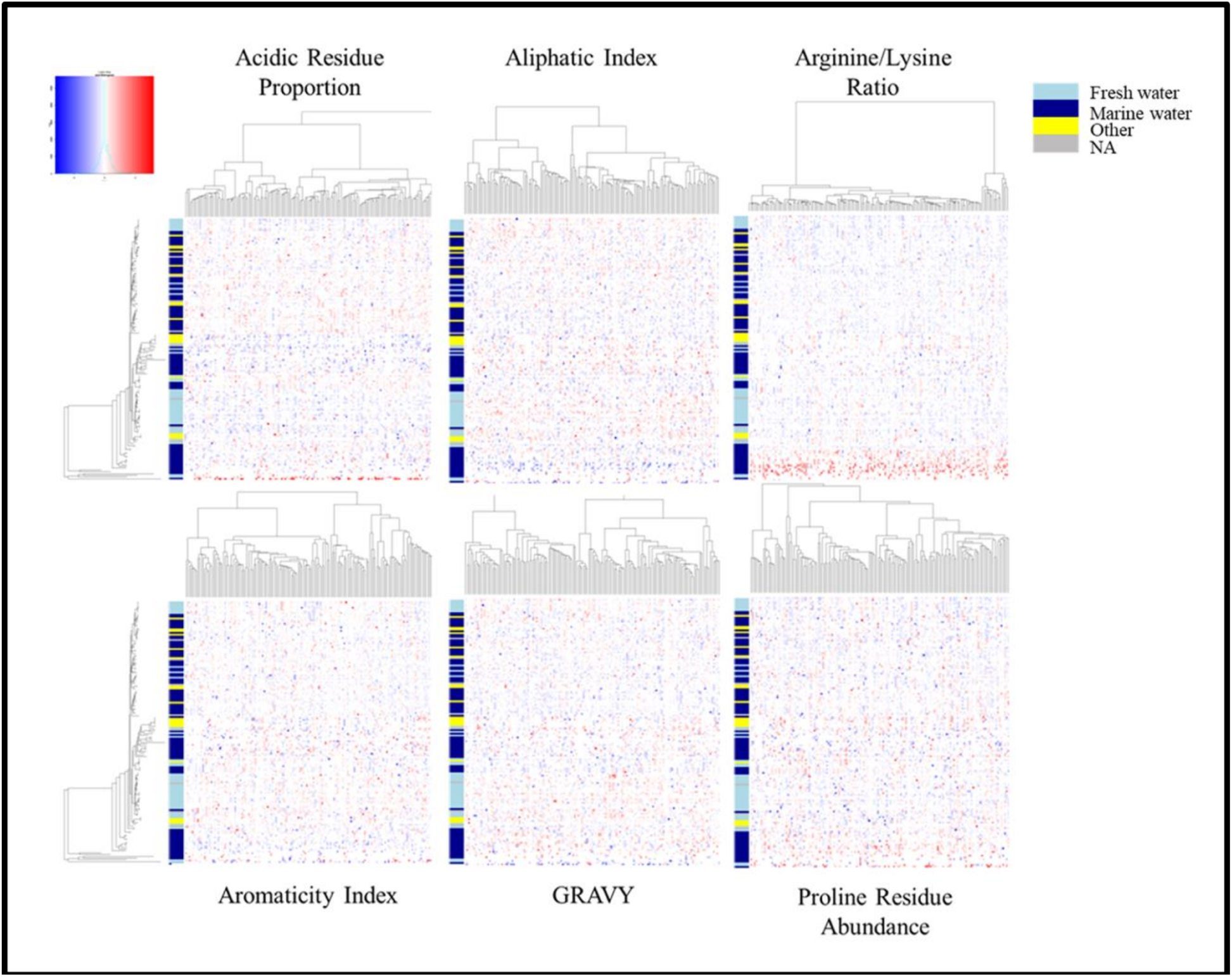
Heatmaps displaying the standardized amino acid compositional indices for the MG-I MAGs (y-axis; phylogenetic distance of each sequence) for each global protein family (x-axis; ward distances). Red indicates a high Z-score, while blue indicates a low Z-score. White indicates a score of zero or an NA. Coloured bars on the left-hand side of each plot indicate the source of each genome (fresh water (light blue); marine water (dark blue); other (yellow); NA (grey)). The outgroup taxa are at the bottom.

**Figure 4:**
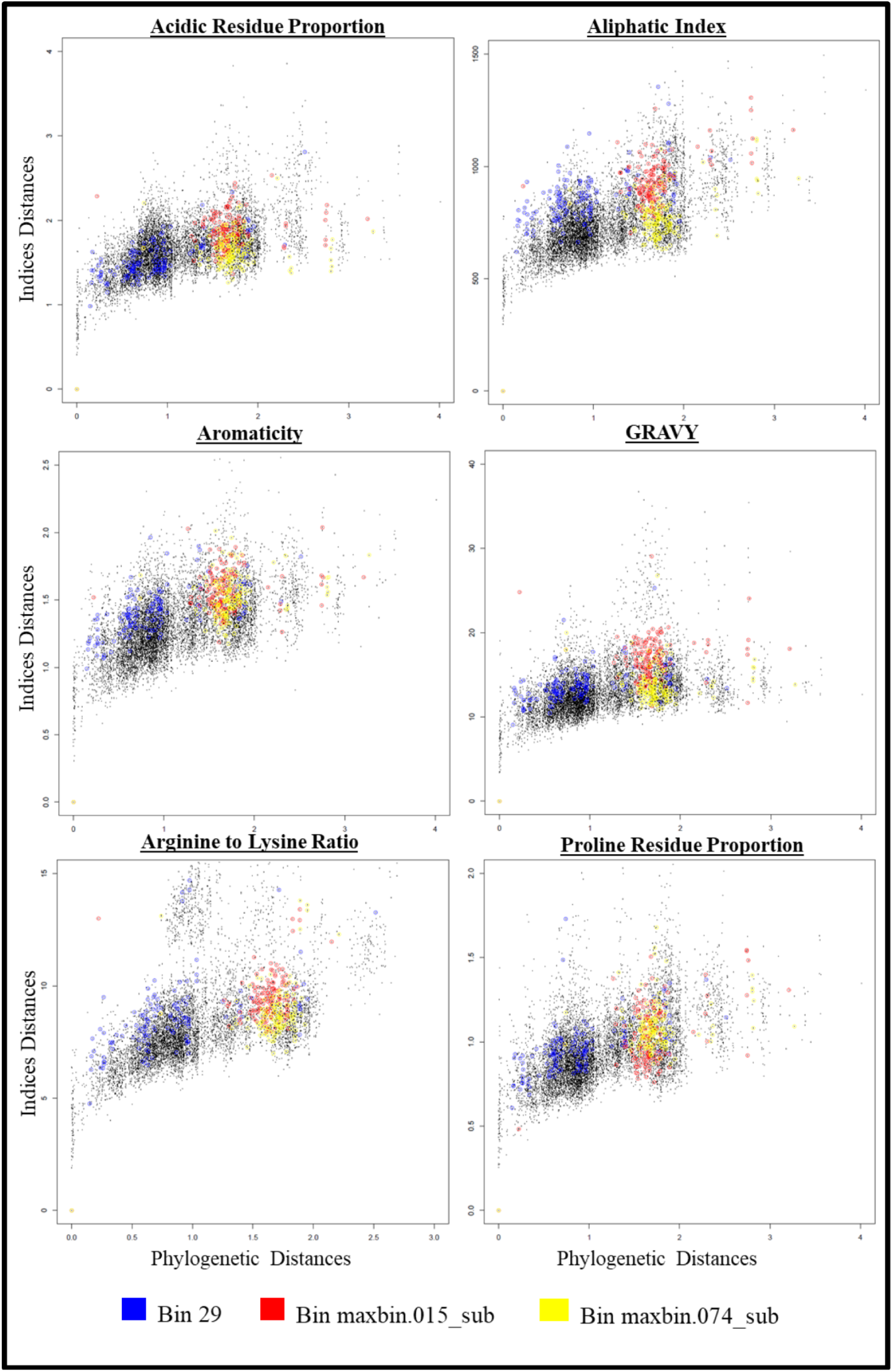
Indices distances against the phylogenetic distances for MG-I highlighting the Western Arctic Archaea. Three MG-I MAGs from the Western Arctic Ocean (Green and Collins, 2026) were included in this analysis.

**Figure 5:**
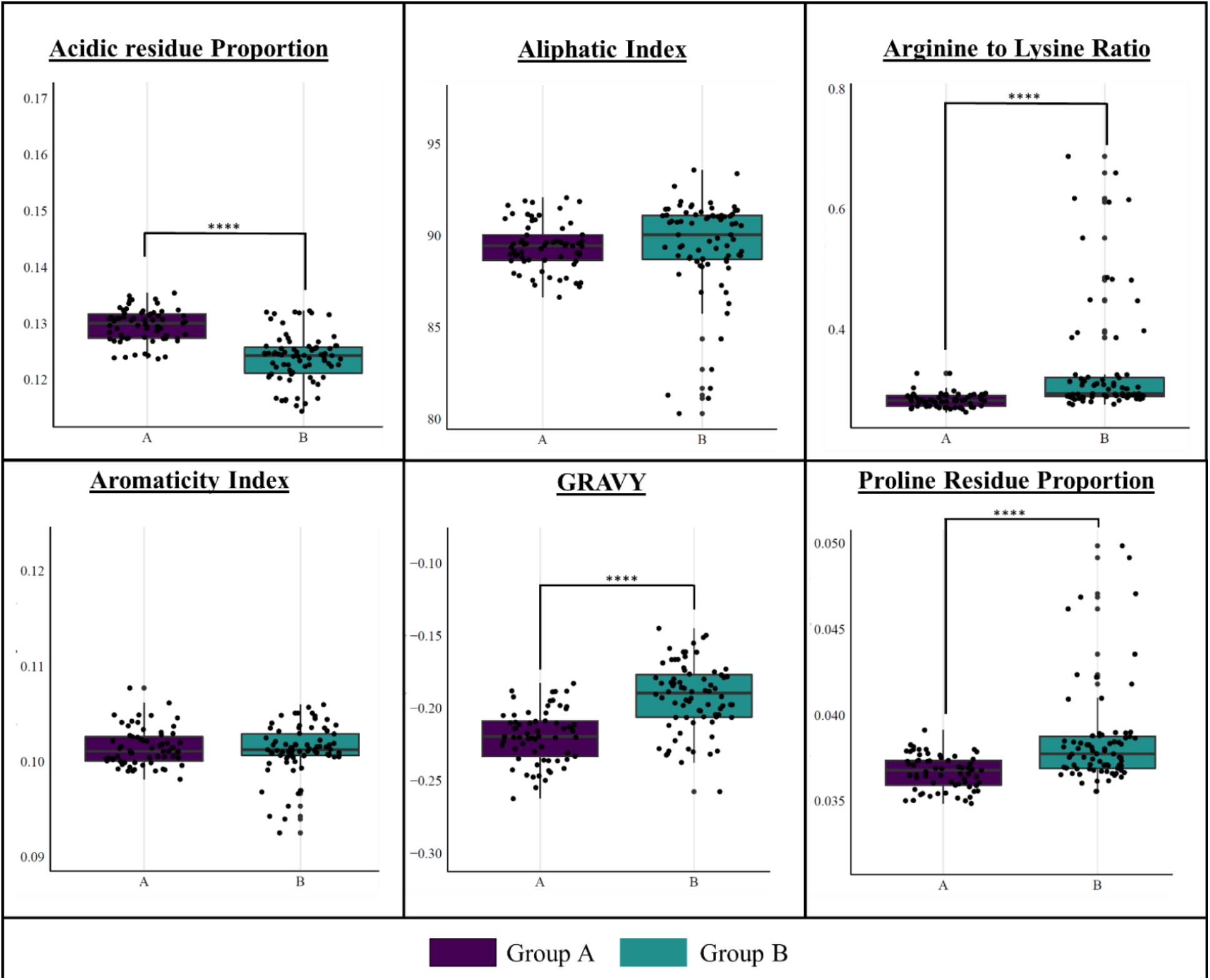
Comparison of indices from the Group A (purple) and Group B (teal) MG-I.

**Figure 6:**
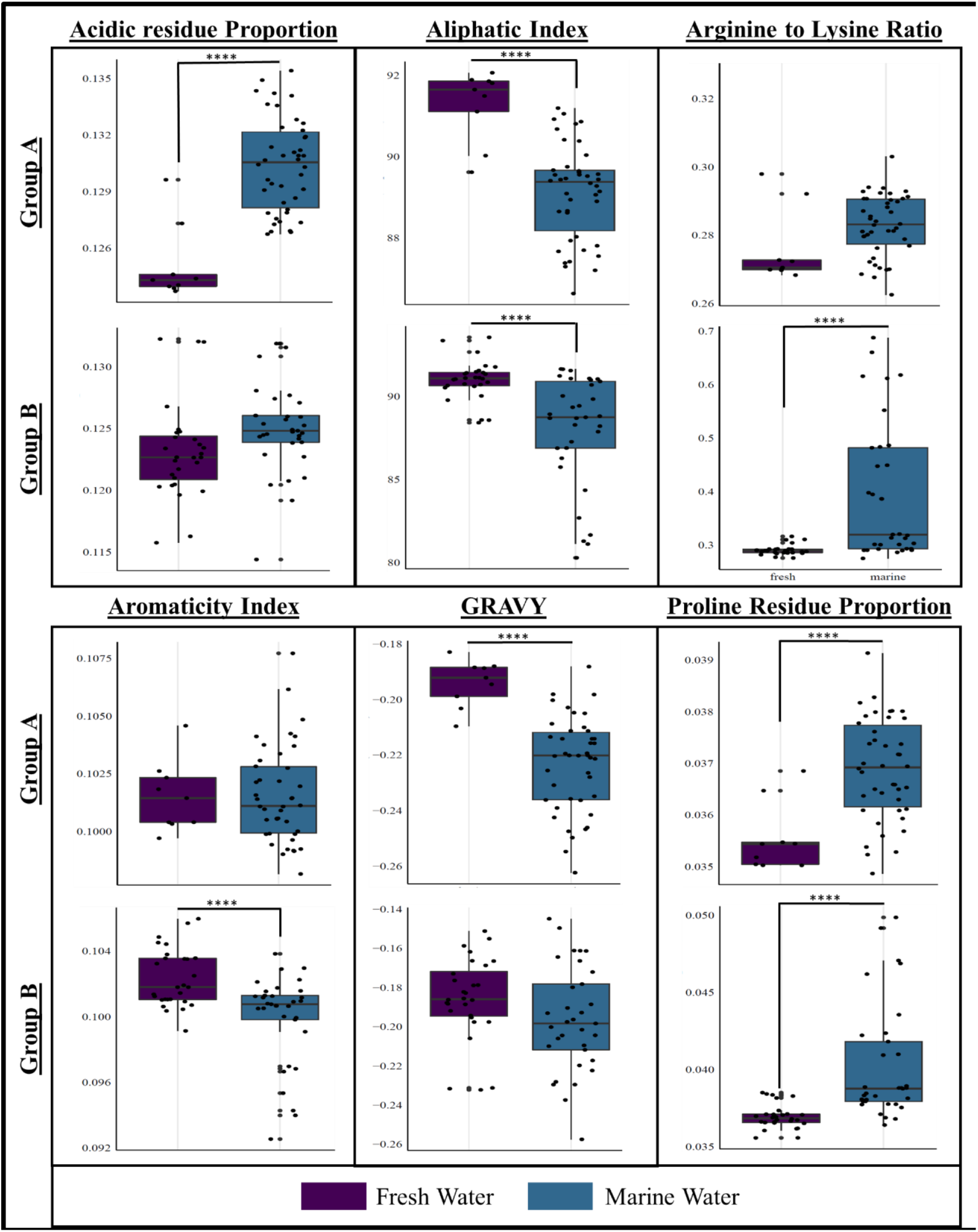
Boxplot of the comparison between fresh water (purple) and marine water (blue) for Group A and Group B MG-I for the amino acid indices.

### 3.2 Euryarchaeota genomes form five clusters based on amino acid compositional indices

Using a collection of 146 Euryarchaeota genomes collected from the BV-BRC portal (Figure 7), the same analysis was performed to determine if the lack of variance in the amino acid usage was consistent across a larger set of Archaea or was specific to MG-I. The outgroups used in this analysis were *Nitrosopumilus* sp. UBA526, *Nitrosopumilus* sp. UBA241, *Candidatus* Woesearchaeota CG07, and *Candidatus* Woesearchaeota CG08. Contrary to the results for MG-I, Euryarchaeota appeared to have a consistent clustering pattern of five clades across all indices measured (Figure 8). This clustering trend appeared to be consistent with the phylogenetic grouping of the organisms (Supplementary Figure S3), rather than the source of their isolation. The first and largest cluster (Group A) was monophyletic and contained members of the class Halobacteria. This group showed higher arginine to lysine ratios (0.642 *±* 0.182) than the other groups (Table 1). The second cluster (Group B) was monophyletic and contained members from the order Methanomicrobiales. This group displayed higher aliphatic indices (92.106 *±* 3.328), and lower arginine to lysine ratios (0.364 *±* 0.172) and proline residue proportions (0.036 *±* 0.005). The third cluster (Group C) was monophyletic and contained members from the orders Methanosarcinales and Methanotrichales. This group showed higher aliphatic (88.55 *±* 4.23) and GRAVY indices (−0.138 *±* 0.075). The fourth cluster (Group D) was polyphyletic and contained members of the orders Archaeoglobales and Methanobacteriales. This group showed lower arginine to lysine ratios (0.595 *±* 0.191), as well as higher GRAVY (−0.137 *±* 0.081) and aliphatic indices (89.006 *±* 5.130). The fifth cluster (Group E) was monophyletic and contained members from the order Thermococcales. This group showed higher average arginine to lysine ratios (0.700 *±* 0.147) and lower average GRAVY index (−0.207 *±* 0.067).

**Figure 7:**
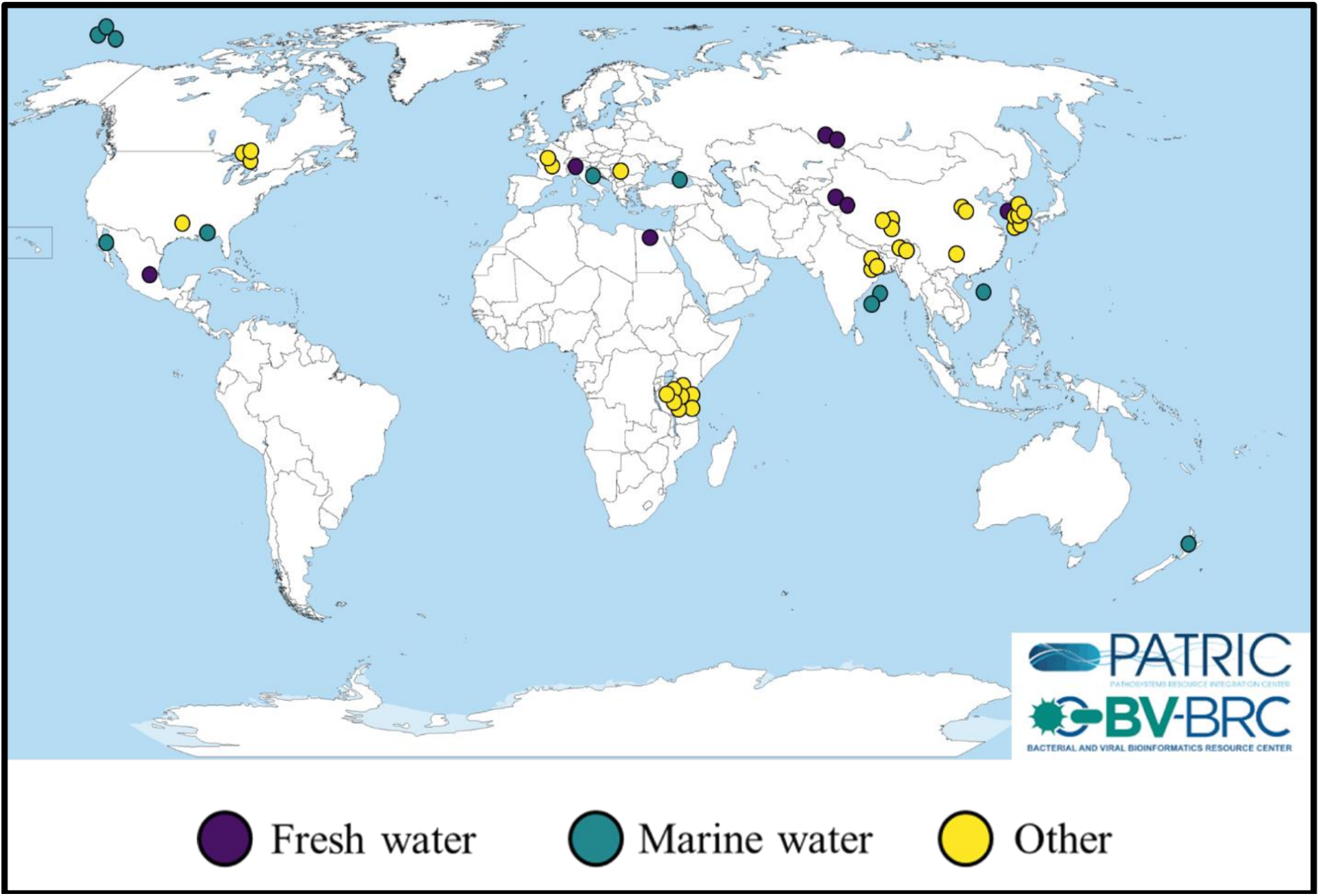
Geographical map showing the sampling locations of the Euryarchaeota genomes obtained from the BV-BRC database. Colours of each circle indicate the sampling source: fresh water (purple), marine water (teal), and other (yellow).

**Figure 8:**
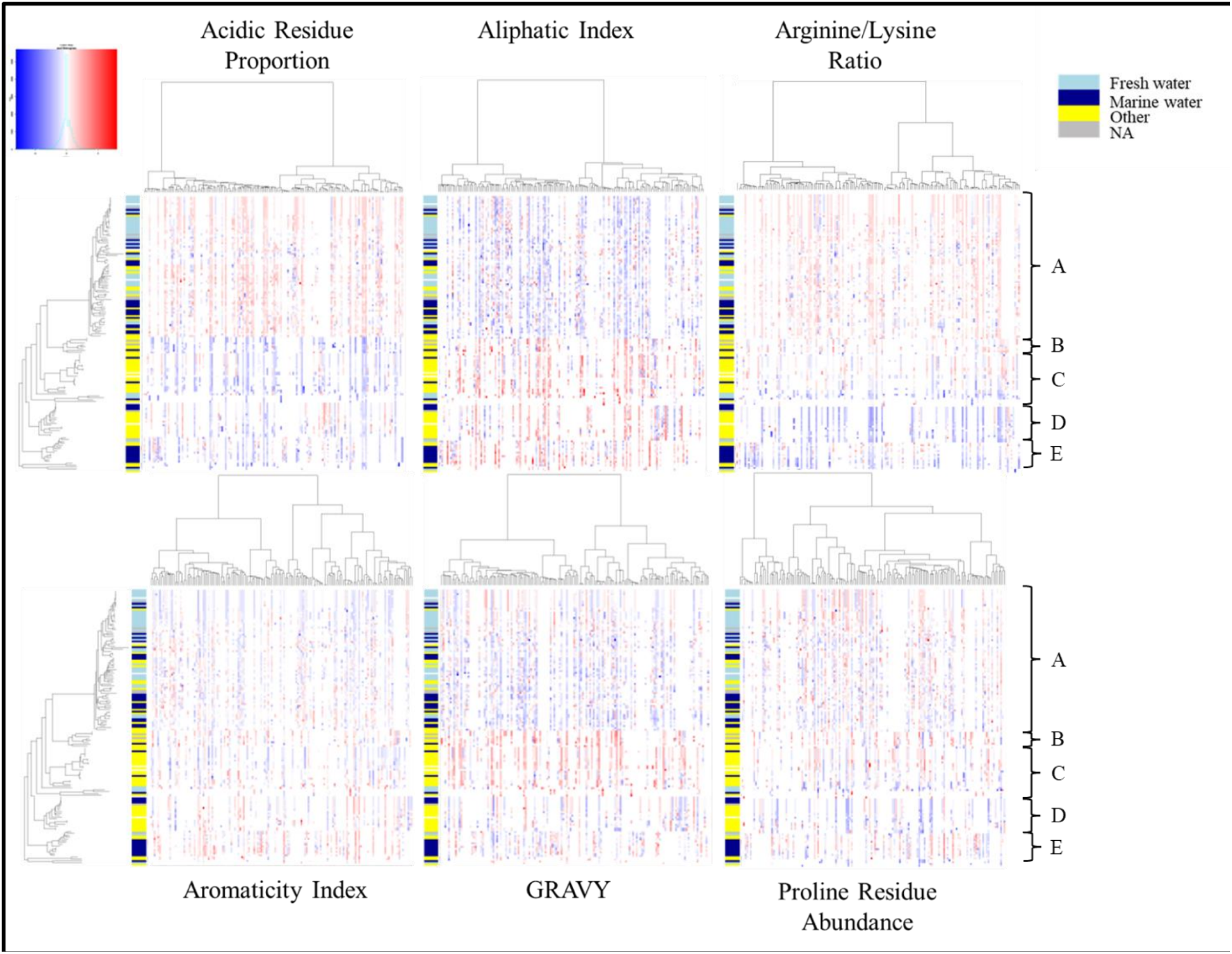
Heatmaps displaying the standardized amino acid compositional indices for the Euryarchaeota genomes/MAG (y-axis; phylogenetic distance of each sequence) for each global protein family (x-axis; ward distance). Red indicates a high Z-score, while blue indicates a low Z-score. White indicates a score of zero or an NA. Coloured bars on the left-hand side of each plot indicate the source of each genome (fresh water (light blue); marine water (dark blue); other (yellow); NA (grey)). The letters on the right-hand side of the plots indicate the clustering. The outgroup taxa are at the bottom.

**Table 1:** Mean (*±* SD) amino acid compositional indices for the five clusters of Euryarchaeota identified in. **Figure 8**.

| Group | Families | Arg/Lys ratio | Acidic residue prop. | GRAVY | Proline residue prop. | Aromaticity | Aliphatic index |
|---|---|---|---|---|---|---|---|
| A | Haloarculaceae, Halobacteriaceae, Halococcaceae, Haloferacaceae, Halorubraceae, Natrialbaceae | 0.645 ± 0.182 | 0.154 ± 0.018 | −0.181 ± 0.056 | 0.045 ± 0.004 | 0.093 ± 0.006 | 86.570 ± 5.060 |
| B | Methanocalculaceae, Methanocorpuscu-laceae, Methanomicrobi-aceae, Methanoregulaceae | 0.364 ± 0.172 | 0.145 ± 0.017 | −0.167 ± 0.046 | 0.036 ± 0.005 | 0.098 ± 0.005 | 92.106 ± 3.628 |
| C | *Candidatus* Methanoperede-naceae, Methanosarcinaceae, Methanotrichaceae | 0.614 ± 0.133 | 0.146 ± 0.019 | −0.138 ± 0.075 | 0.045 ± 0.004 | 0.094 ± 0.005 | 88.55 ± 4.230 |
| D | Archaeoglobaceae | 0.595 ± 0.191 | 0.146 ± 0.021 | −0.137 ± 0.081 | 0.043 ± 0.006 | 0.097 ± 0.006 | 89.006 ± 5.130 |
| E | Thermococcaceae | 0.700 ± 0.147 | 0.164 ± 0.017 | −0.207 ± 0.067 | 0.045 ± 0.002 | 0.095 ± 0.008 | 85.375 ± 5.760 |
| Outgroup | NA | 0.675 ± 0.171 | 0.157 ± 0.016 | −0.157 ± 0.060 | 0.044 ± 0.002 | 0.096 ± 0.008 | 87.467 ± 6.965 |

### 3.3 Sponge-associated MG-I MAGs differ in arginine to lysine ratio, amino acid usage, and codon usage

In investigating the amino acid compositional indices for cold adaptation, one specific group of MG-I was found to have unusually high arginine to lysine ratios (Figure 9A). The MAGs in this group were all found to have been isolated from within or on sea sponges (Table 2). These genomes were all members of the Nitrosopumilaceae family, and all originated from marine environments. Additionally, this set of MAGs, which will be referred to as sponge-associated MAGs (SAMs) for the remainder of the text, were also previously identified as members of the outlier *Nitrosopumilus* MAGs that were incorrectly phylogenetically clustering with the Group B MG-I (Section 3.1). Not all of these genomes were SAMs, and not all the SAMs were grouped here. The SAMs included in this list were Nitrosopumilaceae archaeon strain Plut 88866, Nitrosopumilaceae archaeon strain Plut 88877, Nitrosopumilaceae archaeon strain PNGco C binSS26, *Nitrosopumilus* sp. strain PNGco C binVS21, *Nitrosopumilus* sp. strain PNGco S binVS9, *Nitrosopumilus* sp. H8, *Nitrosopumilus* sp. D6, *Nitrosopumilus* sp. H13, *Nitrosopumilus* sp. B06, Nitrosopumilaceae archaeon p16x60bins.31, Nitrosopumilaceae archaeon p16x44bins.32, Nitrosopumilaceae archaeon 277c 17, and Nitrosopumilaceae archaeon 277c 18. Figure 9B shows that the arginine to lysine ratio indices for the SAMs were significantly higher (*P <* 0.001) than MG-I found in other environments (*n*_1_ = 19 (SAMs); *n*_2_ = 128 (fresh, marine, and other), degrees of freedom = 145); however, the variance was greater in the non-SAMs group. Significant differences were also observed in the aliphatic (*P <* 0.00001), aromaticity (*P <* 0.05), and proline residue proportion (*P <* 0.000001) indices. In Figure 10, it is seen that the patterns of amino acid usage by MG-I are consistent across the majority of the genomes measured. The SAMs, however, showed a differential pattern of amino acid utilization. The only amino acids that did not show a significant difference in the SAMs were alanine and lysine (Figure 11). Amino acids used less frequently in the SAMs were tyrosine, serine, arginine, proline, asparagine, methionine, leucine, phenylalanine, glutamic acid, and aspartic acid, while the amino acids used more frequently were tryptophan, valine, threonine, glutamine, isoleucine, histidine, glycine, and cysteine. Acidic amino acids, like glutamic acid and aspartic acid were used less frequently in these organisms which was consistent with the averaged indices for the acidic residue proportion (0.125 in SAMs and 0.127 in MG-I), although this difference was not significant (*P >* 0.05). The contradiction between the acidic residue proportions and the individual glutamic acid and aspartic acid utilization frequencies was due to the nature of the calculations. The index calculates a total count of acidic amino acids in the proteins of each genome, while the frequency calculation takes into consideration the length of the protein sequences, therefore, removing the bias of different genome sizes. Bulky aromatic amino acids like tyrosine and phenylalanine were utilized less frequently, however within the same class of amino acid, tryptophan was used more frequently. The aromaticity index was consistent with these results, being lower in the SAMs than in the rest of MG-I (*P <* 0.05). Histidine was used more frequently than arginine, both basic amino acids. Valine and isoleucine were also more frequently used, having shorter aliphatic side chains than methionine and leucine, which were not used as frequently; consistent with this, the aliphatic index was significantly different in the SAMs (*P <* 0.00001). Glycine and cysteine were observed more frequently. Alternatively, proline was observed less frequently. Glutamine and asparagine are both polar uncharged amino acids which show differential utilizations. Glutamine being used more frequently, however larger, is coded by the codons CAA and CAG, which have more C/G than the codons encoding asparagine (AAU and AAC). To further explore these differences, codon usage by each group was analyzed. It is known now that these organisms show preferential usage of different amino acids, however, the codons used to encode these amino acids were also different, specifically in the wobble position of the codon. Figure 12 shows that the SAMs differed in their codon usage compared to the MG-I. The SAMs also had higher overall genome GC content (Figure 13A) than the others. Additionally, within the coding regions of the genomes, the SAMs showed a preference towards codons with a higher GC content (Figure 13B).

**Figure 9:**
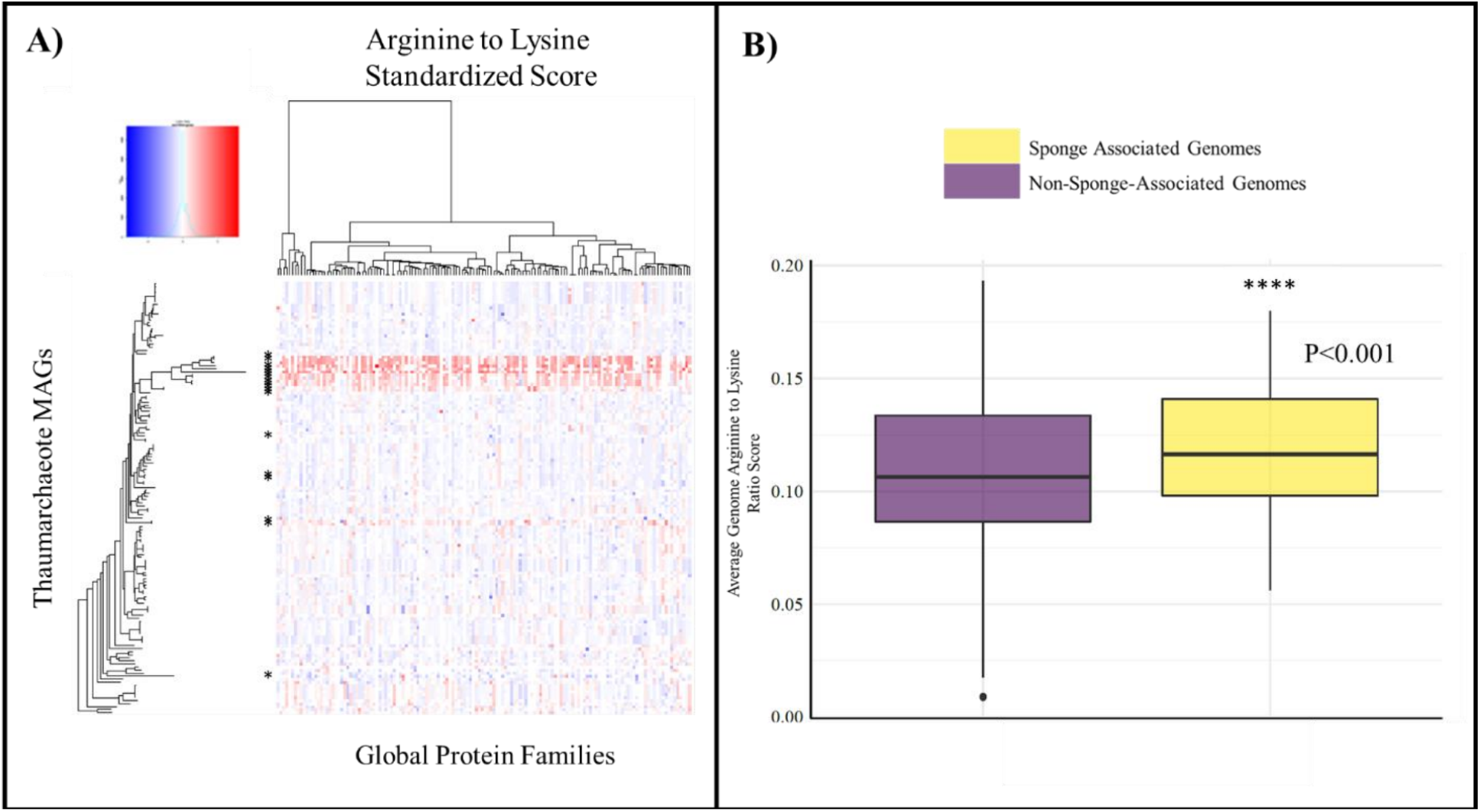
Arginine to lysine Z-scores for the MG-I sponge-associated MAGs. A) Heatmap displaying the standardized arginine to lysine Z-scores for the MG-I genomes/MAGs (y-axis; phylogenetic distance of each sequence) for each global protein family (x-axis; ward distance). Red indicates a high score, while blue indicates a low Z-score. White indicates a Z-score of zero or an NA. An asterisk (*) indicates a sponge-associated genome. B) Boxplot of the average arginine to lysine ratios of the sponge-associated MAGs compared to the other MG-I genomes.

**Figure 10:**
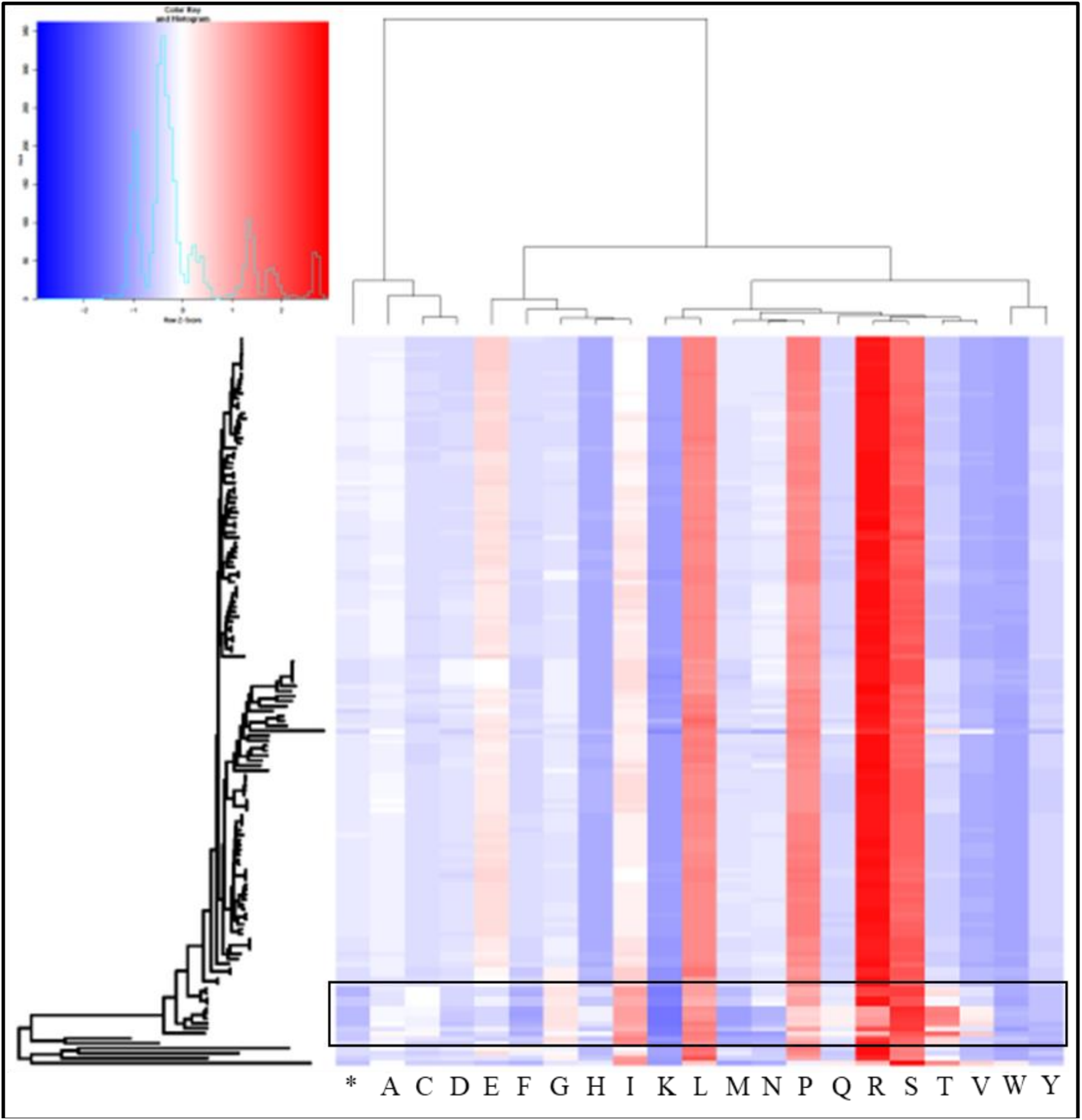
Heatmap displaying the standardized frequencies of the amino acids (x-axis) used in the MG-I MAGs (y-axis). Blue indicates lower usage and red indicates higher usage.

**Figure 11:**
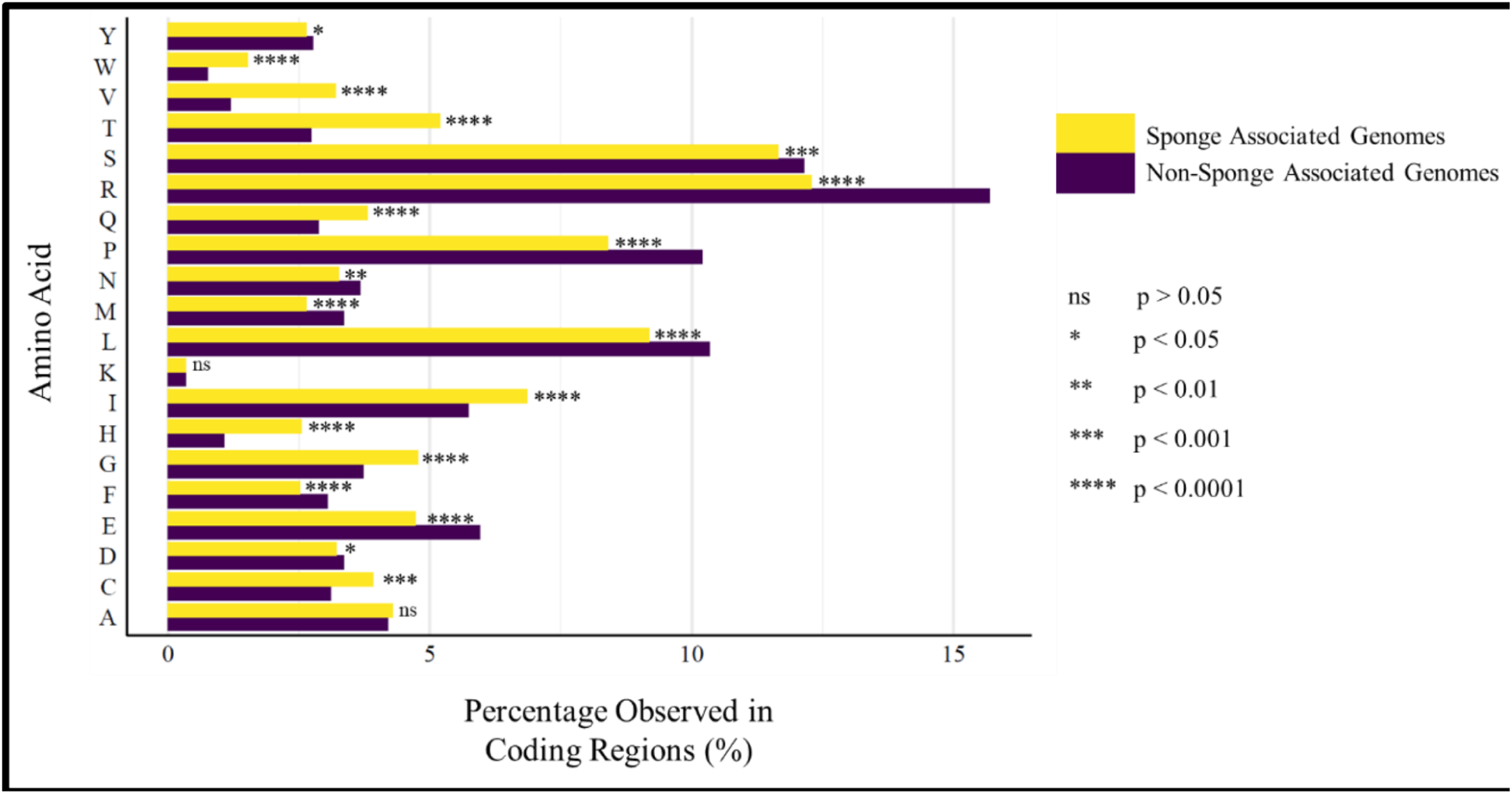
Bar plot displaying the percentage of each amino acid observed in the genomes of the sponge-associated MAGs and non-sponge associated MAGs.

**Figure 12:**
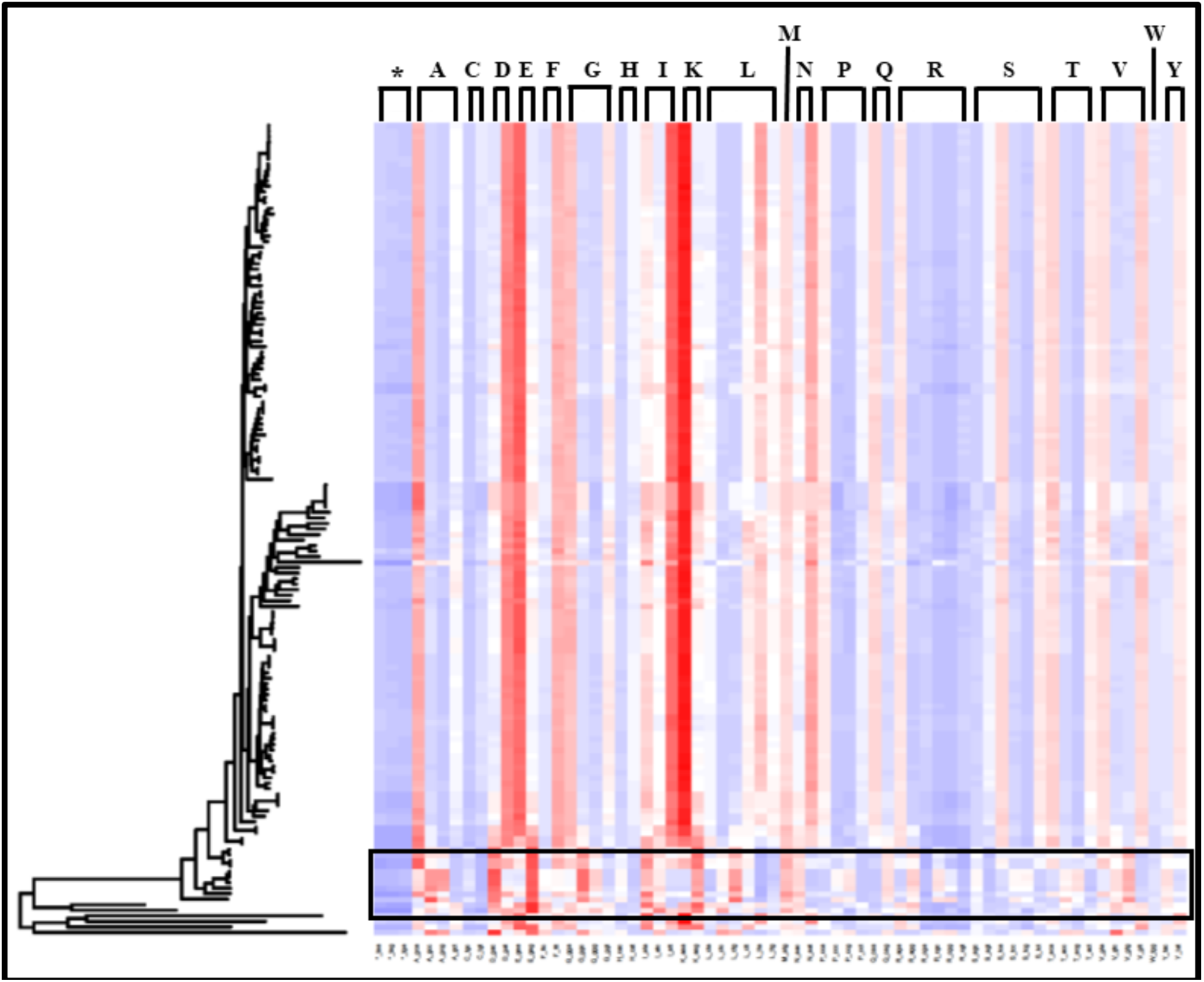
Heatmap of codon usage in MG-I. Y-axis is the phylogenetic tree of the MG-I genomes. X-axis are the codons differentiated by the amino acids these encode. The amino acids are listed at the top as their single letter code. The black box highlights the sponge-associated genomes.

**Figure 13:**
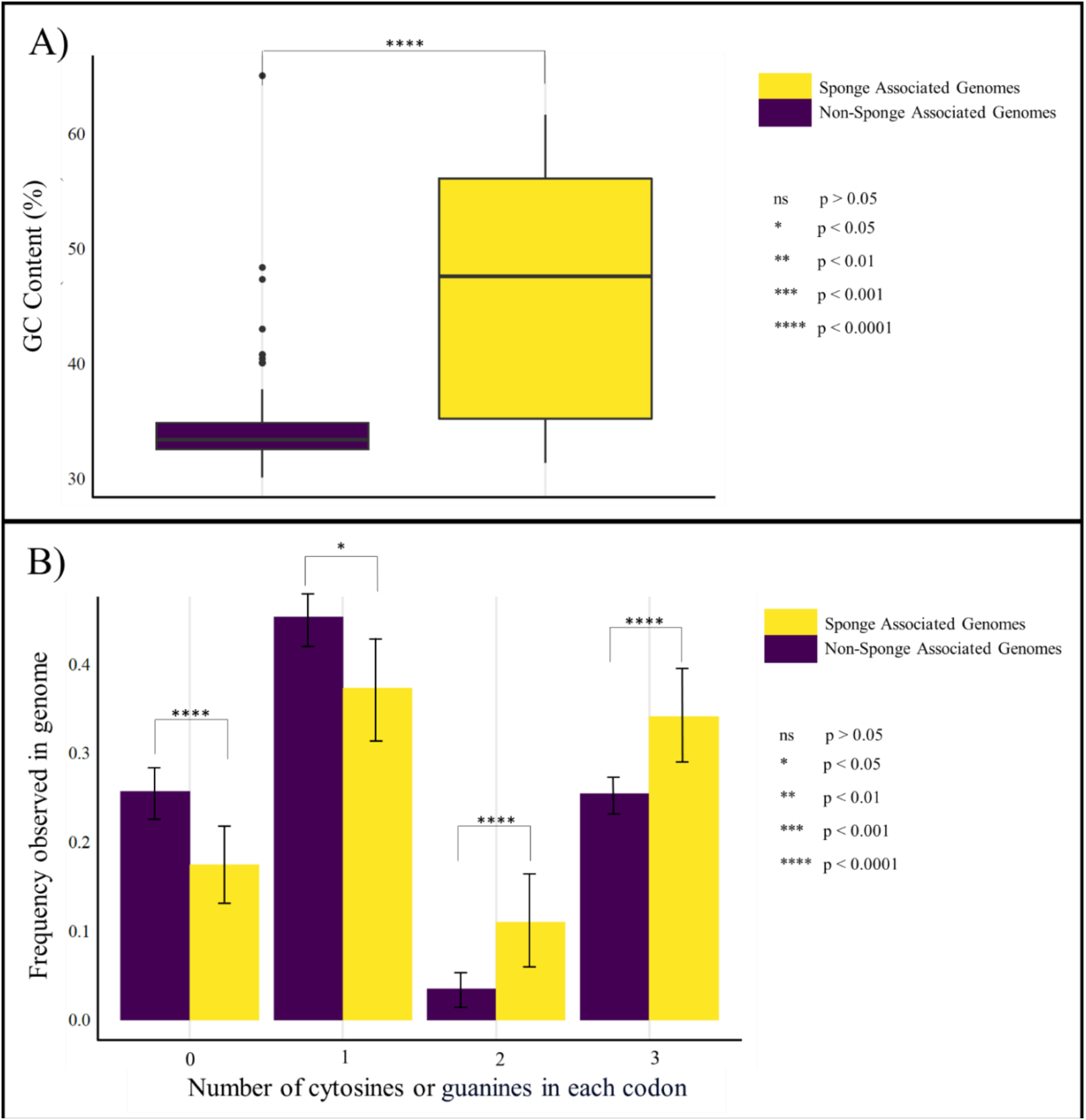
GC content of the sponge-associated MAGs compared to the other members of MG-I: A) overall genomic GC content. B) the frequency of codons observed in the MAGs which have either zero, one, two or three guanines or cytosines.

**Table 2:** Sponge-associated MG-I MAGs (SAMs), their host, and their location of origin.

| Genome name | Host name | Location |
|---|---|---|
| *Nitrosopumilus* sp. Nsub | *Suberites* | NA |
| *Nitrosopumilus* sp. strain HM N | *Hymedesmia methanophila* | Mexico |
| Nitrosopumilaceae archaeon strain Plut 88866 | *Porites lutea* | Australia |
| Nitrosopumilaceae archaeon strain Plut 88877 | *Porites lutea* | Australia |
| Nitrosopumilaceae archaeon strain PNGco C binSS26 | *Coelocarteria singaporensis* | Papua New Guinea |
| *Nitrosopumilus* sp. strain PNGco C binVS21 | *Coelocarteria singaporensis* | Papua New Guinea |
| *Nitrosopumilus* sp. strain PNGco S binVS9 | *Coelocarteria singaporensis* | Papua New Guinea |
| *Nitrosopumilus* sp. H8 | *Hexadella dendritifera* | Atlantic Ocean |
| *Nitrosopumilus* sp. D6 | *Hexadella dendritifera* | Atlantic Ocean |
| *Nitrosopumilus* sp. H13 | *Hexadella dendritifera* | Atlantic Ocean |
| *Nitrosopumilus* sp. B06 | *Hexadella dendritifera* | United Kingdom |
| Nitrosopumilaceae archaeon p16x60bins.31 | *Ircinia* sp. | Panama |
| Nitrosopumilaceae archaeon p16x44bins.32 | *Ircinia* sp. | Panama |
| Nitrosopumilaceae archaeon 277c 17 | *Petrosia ficiformis* | Israel |
| Nitrosopumilaceae archaeon 277c 18 | *Petrosia ficiformis* | Israel |
| Nitrosopumilaceae archaeon 277c 19 | *Petrosia ficiformis* | Israel |
| *Nitrosopumilus* sp. strain PfA | *Petrosia ficiformis* | Israel |
| *Nitrosopumilus* sp. strain MH-Pat-all metabat2 16 | *Mycale hentscheli* | New Zealand |
| *Nitrosopumilus* sp. LS AOA | *Lophophysema* sp. RT-2015 | NA |

### 3.4 MG-I phylogeny is not consistent with gene content

MG-I evolution has been a continuing topic throughout this study. It was observed that there were very few differences in the six amino acid index measures and that expected trends for phylogeny were not followed. The expected trends for the phylogenetic and environmental niches displayed varied patterns for the indices, although not associated with cold adaptation. In light of these variations, the presence/absence of archaeal global protein families (GPFs) was analyzed to determine changes in proteomic composition that may affect cold adaptation (Figure 14). The MAGs grouped into two large clusters, with the latter dominated by lower quality MAGs. This latter group also had a greater proportion of absent gene families. Associated genomes with similar gene content could be clustered together because they either share a common ancestor or because the genomes are not complete. The relationship between the phylogenetic grouping of the MAGs and distance measurements of the global protein family presence or absence data was also analyzed (Figure 15; Supplementary Figure S4), showing that the phylogenetic grouping is not consistent with the distance measurements. Within the cluster of good-quality MAGs, however, the dendrogram structure largely followed phylogeny. The index distances and phylogenetic distances between all of the MG-I MAGs were analyzed (Figure 16). The outgroups (one representative is highlighted in grey) are the most phylogenetically distant, as expected. One of the SAMs was chosen (2202732.34) as a representative to display that they were more phylogenetically related to the MG-I than the outgroups, but they generally were the most phylogenetically distant from the main population. The arginine to lysine ratio distances for the SAMs were distinct, as many of them clustered in the top left signifying a greater distance in the index compared to other MG-I at similar phylogenetic distances. In Section 3.1 (Figure 4; a zoomed-in portion of the same figure) it was observed that Western Arctic archaeal representatives were closer to the general MG-I population in both index distances and phylogenetic distances.

**Figure 14:**
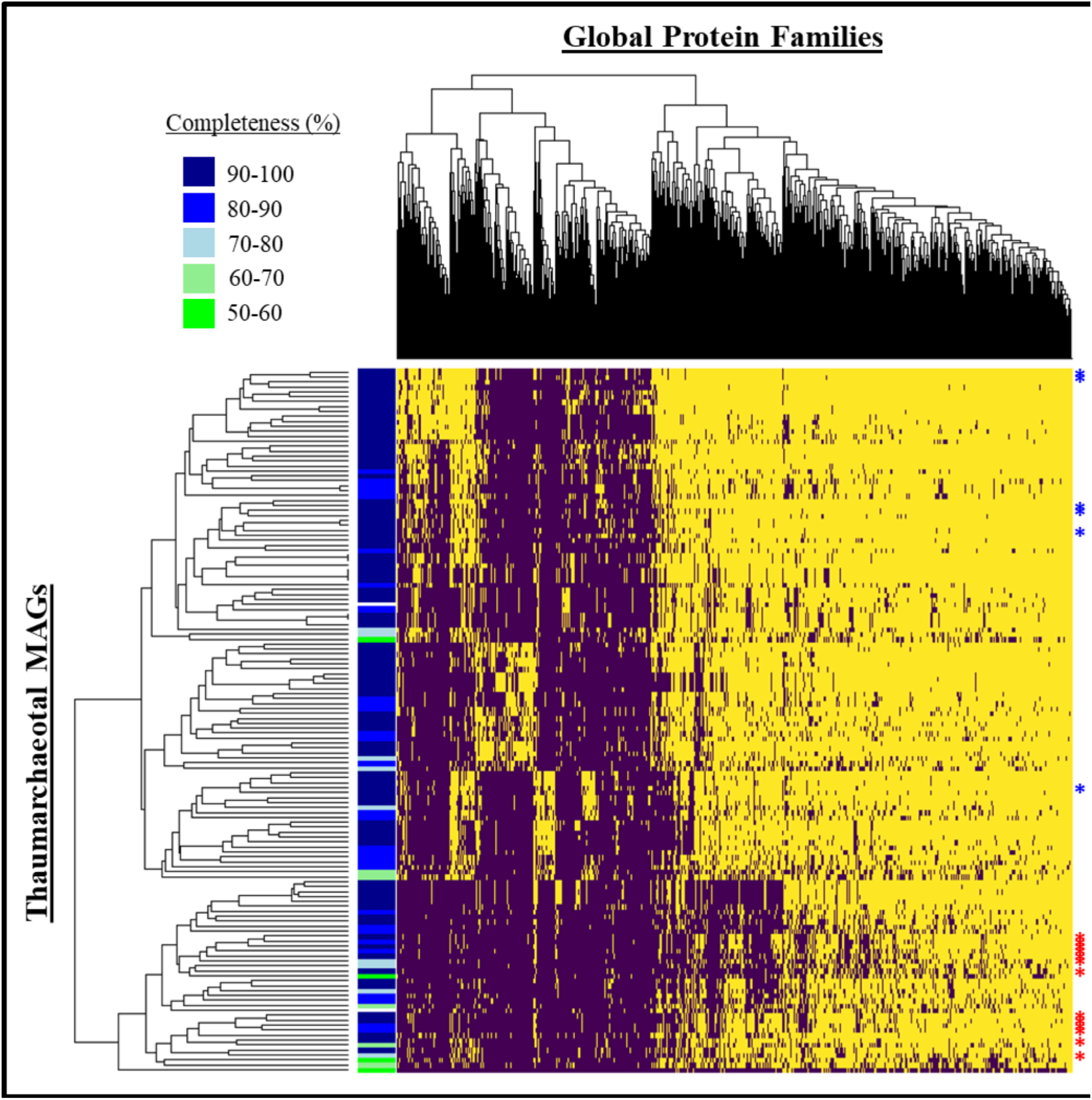
Presence/absence heatmap for the MG-I genomes. Yellow indicates presence and purple indicates absence. Coloured bar on the left indicates completeness. Sponge-associated MAGs (SAMs) belonging to MG-I phylogenetic group A (blue) and B (red) are identified by asterisks (*) in the right-hand side.

**Figure 15:**
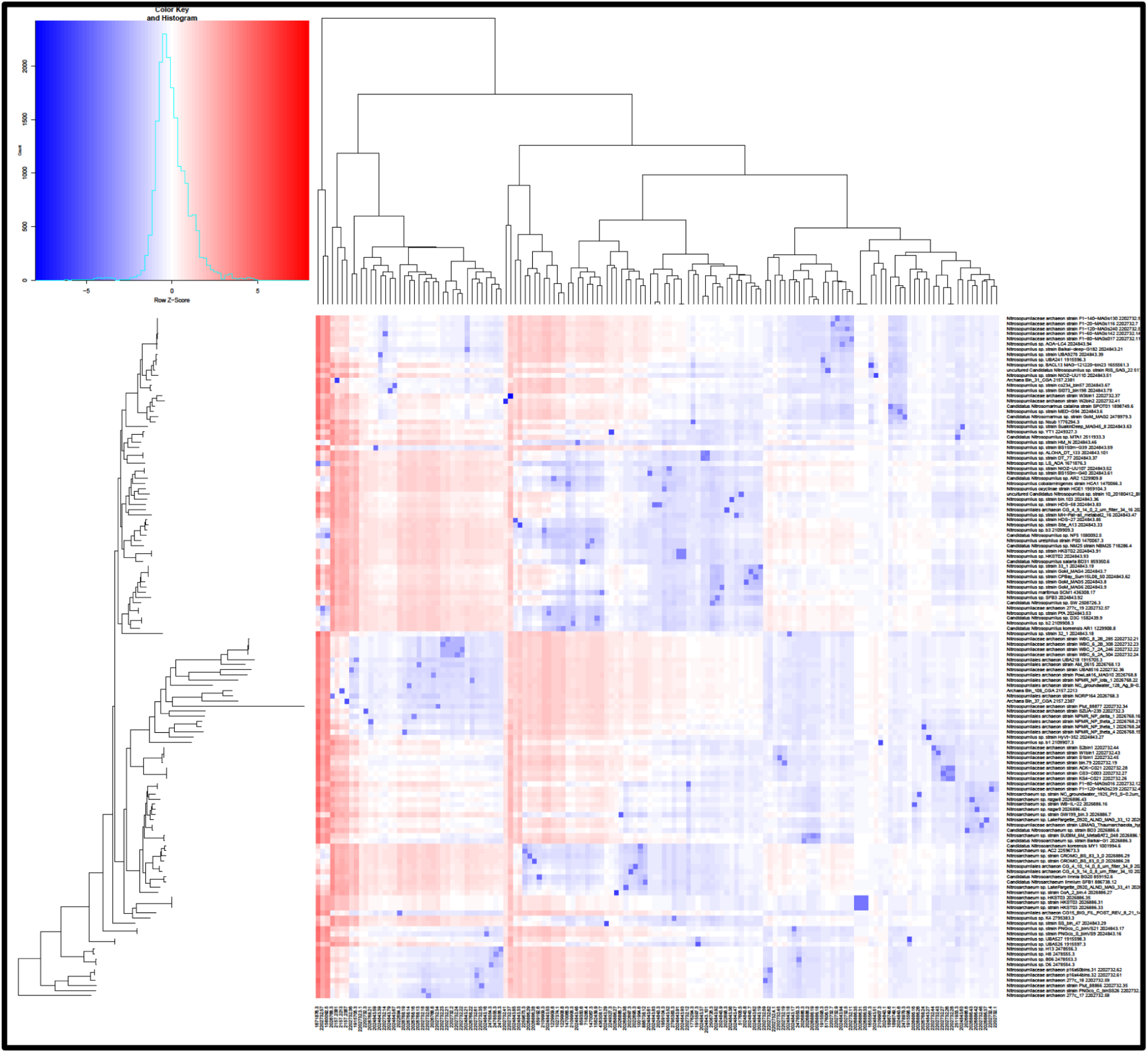
Heat map of phylogeny (y-axis) and presence/absence of global protein distances (x-axis) for the MG-I. Blue indicates a lower distance and red indicates a larger distance between MAGs. Dark blue indicates a direct comparison between the same MAG.

**Figure 16:**
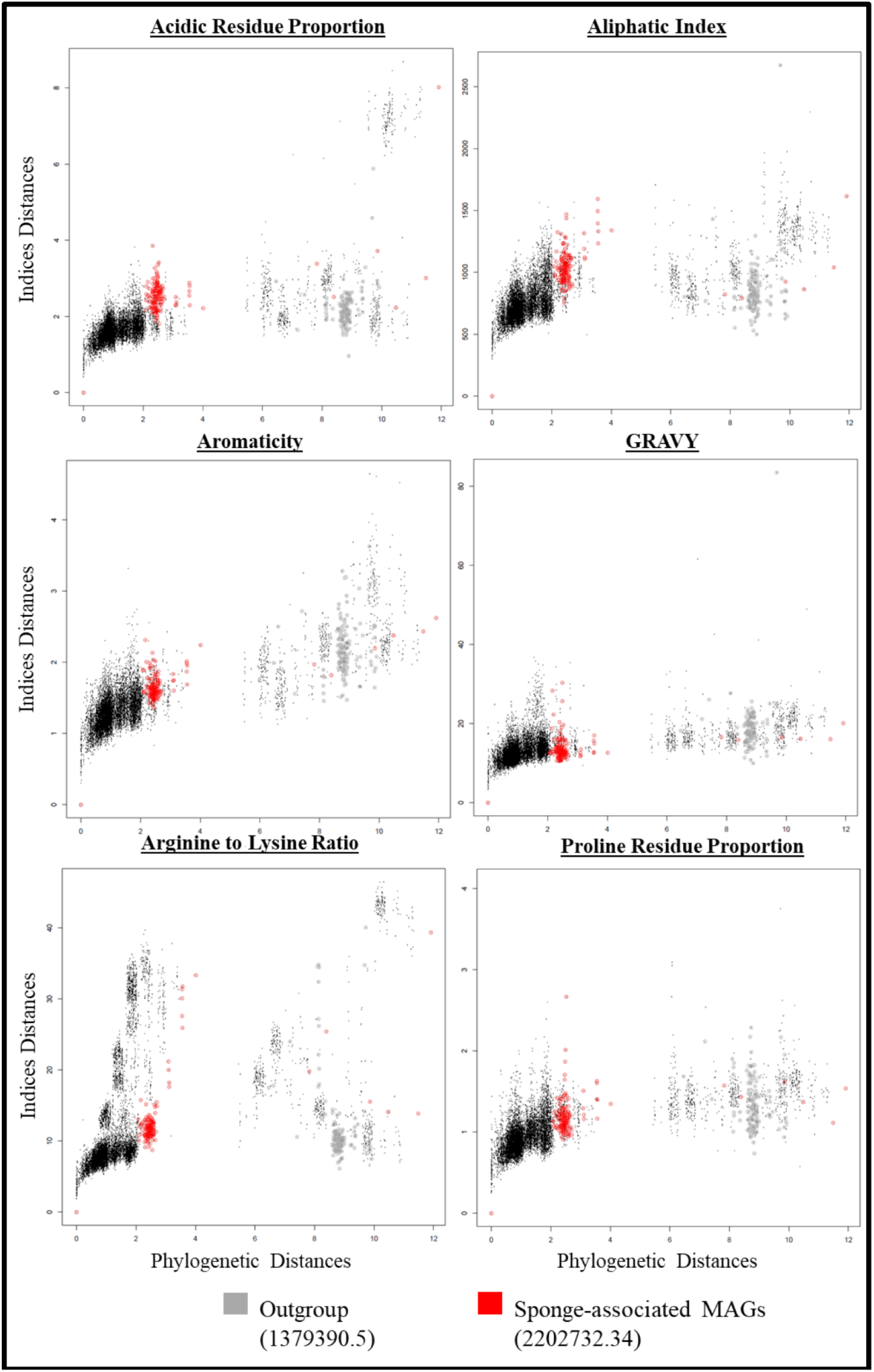
Indices distances against the phylogenetic distances for MG-I. One SAM (genome ID: 2202732.34) is highlighted here alongside one of the outgroup MAGs (genome ID: 1379390.5).

## 4 Discussion

### 4.1 Efficacy of amino acid compositional indices for assessing cold adaptation in Archaea

Inferred cold adaptation in Archaea was measured using six established amino acid compositional indices: arginine to lysine ratio, acidic residue proportion, GRAVY index, aliphatic index, proline residue proportion, and aromaticity index. The hypotheses were that there would be signatures of temperature-adaptation in the proteomes of marine Archaea and that environmental temperature would be a factor influencing the gene content and amino acid usage in Archaea. Based on analyses performed with Bacteria (Gentilhomme, 2020), it was expected that the amino acid indices would be lower in MAGs from low temperature environments and that the phylogeny would reflect the environmental niches of marine Archaea. Since a large collection of archaeal MAGs was being analyzed, with diverse sampling locations, sources, seasons, and years, it was expected that there would be a high level of variation amongst the MAGs that would be consistent with their phylogenetics and environmental niches. The phylogenetic and amino acid compositional index distances were calculated (Figures 3 and 4) and showed that the Arctic MG-I MAGs (Green and Collins, 2026) were all well within the normal distribution of distances in both the index distances and phylogenetic distances. Since these MAGs were collected from an environment with lower temperatures, it was expected that the distances for the indices would be greater for them compared to the rest of the population. Therefore, it was concluded that these indices did not show apparent signatures of cold adaptation in Archaea. Potentially this speaks to the efficacy of the indices themselves. A review study (Li et al., 2019) that looked at a collection of different genomic and proteomic indices and metrics for inferring optimal growth temperature (OGT) found that although most of these tools are quite good for high temperatures, they do not behave the same way for psychrophiles. For example, GC content is often used as a metric to measure heat-adaptation, where higher GC contents are seen in thermophilic (heat loving) microorganisms. However, in psychrophilic microbes GC content does not correlate with optimal growth temperature (Li et al., 2019). The explanation as to why is unclear. Although the Arctic MG-I MAGs did not show differences based on the amino acid indices tested, there were other parameters that did seem to influence the amino acid usage in this group. The first factor was the phylogenetics. The MG-I organisms were organized phylogenetically into two groups. It was observed that four of the indices tested were significantly different between the groups (Figure 5). It is important to note that the three Arctic MG-I MAGs included in this analysis were found in both of the phylogenetic groups. Additionally, it was determined that these differences were likely a result of the SAMs incorrectly clustering with phylogenetic group B due to long-branch attraction. This will be discussed in the following section. The second factor was the environment or source of collection. The collection sources were categorized as marine water, fresh water, or other. It was shown that there were significant differences in all indices between the marine water and the freshwater MG-I MAGs in at least one of the phylogenetic groups (Figure 6). This observation was consistent within the two phylogenetic groups as the aliphatic (*P <* 0.0001), acidic residue proportion (*P <* 0.0001), GRAVY (*P <* 0.0001), and proline residue proportion (*P <* 0.0001) indices were significantly different in Group A, and the arginine to lysine ratio (*P <* 0.0001), aliphatic (*P <* 0.0001), aromaticity (*P <* 0.0001), and proline residue proportion (*P <* 0.0001) indices were significantly different within Group B. This suggests that the environmental conditions may be correlated with the amino acid composition seen in MG-I. This observation also implies that there is possible convergent evolution occurring where organisms from different lineages are obtaining similar evolutionary traits based on their environments. It was observed that the Euryarchaeota and the MG-I displayed very different patterns of amino acid usage. While there was little variation between the MG-I clades after the removal of the SAMs, the Euryarchaeota had very distinct patterns based on five phylogenetic groups. As an example, the first and largest cluster (Group A) which contained members of the class Halobacteria displayed higher arginine to lysine ratios (0.642 *±* 0.182) than the other groups (Table 1). All members of this class are halophilic (salt-loving) and are known to have generally higher arginine to lysine ratios (Fukuchi et al., 2003) likely due to the protein stability required in a highly saline environment. While there were differences between the different clades of Euryarchaeota, there were very few differences within the MG-I. Recently, it has been proposed that the Euryarchaeota should actually be reclassified from one phylum into five different phyla (Rinke et al., 2020). This new classification was consistent with the phylogenetic grouping seen here. The analysis performed on the Euryarchaeota was consequently performed at the superphylum level, rather than at the phylum level, following this new classification. Rinke et al. (2020) also suggested that the Thaumarchaeota (MG-I), which are members of the TACK superphylum, should be reclassified. They further suggested that the entire TACK superphylum should be combined to create one phylum. This also changes the context in which these results are understood. The resolution at which these indices remain useful may be at a higher taxonomic level in Archaea. These indices have also been tested in Bacteria, such as Colwelliaceae, and were found to show distinctive differences within the family (Gentilhomme, 2020). In that analysis, DNA from isolated strains of Colwelliaceae was sequenced and analyzed in comparison to their OGT. It was found that the amino acid compositional indices tested were different within the different clades of Colwelliaceae. This is inconsistent with the archaeal results even with a much higher phylogenetic resolution in the Colwelliaceae study. In that study, the indices tested were all correlated with OGT, however, not all were positively correlated, as was expected (Gentilhomme, 2020). Aliphatic and GRAVY indices were negatively correlated with OGT in Colwelliaceae. These indices are treated as independent measures in this analysis although the underpinning molecular processes are interconnected. The negative correlations indicate the presence of more hydrophobic amino acids in these genomes, driven by a greater proportion of short-chained hydrophobic amino acids. Additionally, only three of the six indices were moderately or strongly associated with OGT (GRAVY, proline residue proportion, and aliphatic index). Although these indices were useful in determining the proteomic characteristics within the different clades of Colwelliaceae, they continued to lack efficacy in determining genomic cold adaptation characteristics in Archaea. They may not be the most useful indices for determining cold adaptation because they did not take into consideration the interconnections between the indices themselves as well as other aspects of the genomes that are influencing cold adaptation. There are other components within the cells that provide cold tolerance or cold adaptation that were not yet discussed here. These include alterations in signal transduction, nucleotide-, co-enzyme-, and amino acid metabolism, defense mechanisms, energy production and conversion, DNA replication and processing, RNA and protein synthesis, cellular division, tRNA modifications, motility, and secretion (Cavicchioli, 2006). As an individual tool, these indices do not capture the entirety of cold adaptation mechanisms in microbes.

### 4.2 Host adaptation in Archaea

During the phylogenetic analysis of the MG-I organisms, it was observed that there was one group of Nitrosopumilaceae that were incorrectly grouping with the Group B MG-I. In phylogenetics, this observation is called long-branch attraction where distantly related lineages are often incorrectly associated with lineages that they are not related to (Susko and Roger, 2021). This occurs when there are enough changes or differences in the genome that they appear similar to another long-branch lineage. Rather than actually being closely related to one another, they are grouped together solely because they have both undergone a high degree of molecular changes. This bias is partially understood and is likely caused by assuming an overly simplistic model of evolution. This explains why some of the SAMs clustered together and separately from their Nitrosopumilaceae relatives. The state changes likely influencing this phylogenetic outcome will be discussed in this section. This same group was also shown to have unusually high arginine to lysine ratios (Figure 9), and it was determined that they had all been isolated from within or on sea sponges. Further analysis showed that there were significant differences in all but two amino acids in the proteomes of the SAMs when compared to the general population (Figure 11). There was an increase in tryptophan which is a very large amino acid with an indole ring which can stabilize protein structures and domains. In contrast, there was a decrease in phenylalanine, which produces rigidity within a protein and reduces flexibility. Histidine was used more frequently than arginine, both basic amino acids. These results suggest a preference for smaller aliphatic side chains and aromatic properties that can increase the conformational stability of the proteins. Valine and isoleucine were also more frequently used, having shorter aliphatic side chains than methionine and leucine which were not used as frequently. Glycine and proline both influence the kinetics of protein folding involving loops (the patternless regions that connect secondary structures to one another, such as beta sheets and alpha helices). Glycine, observed more frequently, is often found to accelerate this process as its small size destabilizes the secondary structure to form a loop. Also due to its smaller size, it has the ability to create bends within the protein allowing for flexibility and movement. Alternatively, proline, which was observed less frequently, disrupts these secondary structures by lacking the ability to form hydrogen bonds creating a rigid break in the secondary structure. An increase in the use of cysteine suggests a push towards more stability with its ability to form disulfide bonds. Overall, there appears to be a selection for increased protein flexibility, while maintaining some level of moderate stability. Additionally, it was seen that the SAMs also had very high GC content in their DNA compared to the general population (Figure 13). Archaeal amino acid composition is generally dominated by a GC bias but can be further influenced by environmental factors (Roy Chowdhury and Dutta, 2012). GC content is often associated with a higher level of DNA stability to prevent denaturation, but also as a possible defense mechanism against predation (Slaby et al., 2017). Microbes often have defense mechanisms to detect foreign DNA from viruses. These mechanisms often work by using the GC content of incoming foreign DNA as a determinant for whether or not to degrade that DNA. Having a higher microbial GC content can allow the cell to differentiate foreign DNA from self DNA. Sponges consist of channels that filter thousands of litres of water daily introducing foreign DNA, viruses, nutrients, and contaminants (Kenny et al., 2020; Slaby et al., 2017). It has been shown in Bacteria that sponge symbionts have greater expression levels of genes associated with defense, amino acid transport and metabolism, recombination, and repair (Slaby et al., 2017). Although archaeal representatives from the Crenarchaeota have been identified with sea sponges, the gene content and expression of these organisms have not been explored. It appears that the differences in amino acid usage and the higher GC content observed in the SAMs are consistent with bacterial symbionts from sea sponges. It is plausible to assume that the other state changes occurring in Bacteria may also be occurring in Archaea in response to sea sponge host adaptation. For example, higher expression of genes involved in recombination are often seen in bacterial symbionts as a mechanism for horizontal gene transfer. Horizontal gene transfer (HGT) is the uptake of DNA via direct cell to cell contact, free DNA in the environment, or viral infection. Microbes can quickly pick up genes from other members in their community that can provide the necessary tools to withstand the conditions. Following this logic, this could explain the long-branch attraction that was observed in the SAMs as a result of horizontally transferred genes contributing to their diversity and rates of evolution. The differences observed in the SAMs’ amino acid usage, codon usage, GC content, and index and phylogenetic distances suggest that the environmental conditions of the host-adaptation SAMs have influenced the evolution of these organisms.

### 4.3 Evolution of MG-I

As previously discussed, the phylogenetics (evolutionary history) of the MG-I in this study were not consistent with the taxonomy (classification or naming system). The amino acid indices tested in the proteomes of the MG-I did not have any distinct trends with regards to the phylogenetic grouping. The relationship between the phylogenetic grouping of the MAGs and distance measurements of the global protein family presence or absence data was also analyzed (Figure 15), showing that the phylogenetic grouping was not consistent with the distance measurements. One explanation for why this may be occurring is that the number of genes shared between highly distant MAGs is low, but they are similar because the shared genes are core genes (e.g., DNA replication machinery and protein synthesis). Another explanation is that the two main clusters in Figure 14 represent two genomic states: one cluster of MAGs sharing a last common ancestor and one that contains MAGs with low completeness. Within the cluster of good-quality MAGs, however, the dendrogram structure largely followed phylogeny. Although it was determined that these indices were not effective in identifying cold adaptation, they did provide insight into the evolutionary status and evolutionary rates of the MG-I. There are two explanations for these observations: that the MG-I are evolving very slowly or are evolving very quickly. The rates of evolution in Archaea in general may be different than Bacteria simply because they harbor different DNA replication machinery. Many of the DNA replication components found in Archaea are more similar to those in eukaryotic cells than those in bacterial cells (Dey et al., 2016; Klinger et al., 2016). If the MG-I are evolving much slower than other microbes in their environment it may mean that they maintain highly conserved genomes because they grow very slowly. *r*-strategists are organisms that live in unstable environments and will typically undergo rapid reproduction in order to stabilize themselves, while *K*-strategists typically live in stable environments, typically found in large populations, and do not need to undergo the same rapid rates of replication. These differences in lifestyle will affect how they structure their proteins. It could also mean that they grow solitarily to the point where they are not competing with other organisms. This second explanation is less likely because there have been strong interactions observed between the MG-I and picoplankton for ammonia competition (Martens-Habbena et al., 2009). If the MG-I growth rates are very low, then it could mean that they do not have time to “catch-up” once in a colder Arctic environment. It could also mean there is not enough pressure to change and that they already harbour the necessary machinery to survive, but not thrive. If the alternative explanation is true, that the MG-I are actually evolving at very high rates, it means that they are obtaining adaptations through other means. Archaea and Bacteria share similar genome sizes that are both gene dense, however a recent study (Kellner et al., 2018) found that bacterial genome size changed in a larger range than in Archaea and that the overall diversity in Archaea was much lower. Ultimately, there may be other factors that are more important for adaptation depending on the microbial community and the environment. Although we did not see evidence of cold adaptation, there may be other environmental or biological factors that are stronger evolutionary influences. Overall, it appears that at a proteomic level these organisms are fairly conserved. One exception to this claim that was observed was with the MAGs associated with sea sponges.

## 5 Conclusions

Although the use of the established amino acid compositional indices for the detection of cold adaptation did not prove to be effective in Archaea, these measures did shed light on the evolutionary history of these organisms in terms of environmental adaptation. It did not appear that the Arctic MAGs had significant changes in their amino acid composition and were very similar to other MG-I organisms found in other, more temperate locations. This could suggest that these organisms are very conserved and do not evolve quickly. It could also suggest that these organisms are simply migrating through the Arctic Ocean. The rates of evolution are slowed here, due to lower temperatures that slow reaction rates and replication times. It was observed that the majority of MG-I MAGs were fairly similar to one another, with the exceptions of those associated with sea sponges, suggesting that host-adaptation provides an evolutionary pressure. Being able to infer preferred growth temperatures of microorganisms is important, especially for uncultured taxa. Although the amino acid compositional indices did not prove to be a useful tool in this manner, there may be other metrics or indices that can be explored. Wang et al. (2019) recently looked at the differences in prevalence of certain protein domains in a large collection of bacterial proteomes to understand their niche adapted lifestyles. This work was done with bacterial halophiles and thermophiles but has not yet been done in psychrophilic Bacteria or Archaea. Since there were differences observed in the former environments, it may be worthwhile to look for domain differences in psychrophiles as well. This analysis could be done in Archaea, however there is little data on archaeal domains. Potentially taking a broader approach that considers more of the connected molecular systems rather than individual indices may be helpful. It has also been discussed that there are many other aspects of cellular biology that are affected by lower temperatures. These include the presence of cold shock proteins, modifications in the lipid membrane, and DNA replication machinery. In this analysis, we looked at cold adaptation from a larger taxa point of view, but it could be useful to further investigate the archaeal taxa from the Western Arctic. One way to accomplish this is to attempt to culture archaeal representatives from this environment and experimentally determine each of their OGTs, which can be used to correlate various indices to OGT. To understand the evolution of the MG-I to a fuller extent, the degree to which horizontal gene transfer influences the genetic makeup of these organisms can be explored. For the sponge associated MAGs, it would be helpful to analyze the genomic composition and understand what it is about the sponge environment that is influencing the evolution of this group. It would also be a good idea to try to sequence and identify viral DNA to determine how much viral infection and predation affects these cells.

## Acknowledgements

This manuscript is adapted from Chapter 3 of the MSc thesis of K. Green (University of Manitoba, 2023). We thank Drs. C. J. Mundy and Jose Luis Rodriguez-Gil for their guidance throughout this work, and Katlin Bowman, Kyle Dilliplaine, Pete Morton, and everyone aboard the USCGC *Healy* during the 2015 GEOTRACES cruise (HLY1502) who was involved in the collection of the metagenomic samples. We thank Anaïs Gentilhomme for her contributions to the analysis of the cold-adaptation amino acid indices, including the index calculation scripts.

## Data availability

All reference genomes and MAGs analyzed are publicly available from the Bacterial and Viral Bioinformatics Resource Center (https://www.bv-brc.org); BV-BRC genome identifiers for the MG-I genomes are listed in Supplementary Table S1. The Western Arctic Ocean MG-I MAGs were reconstructed from metagenomic data available under NCBI/ENA BioProject accessions PRJEB14154 and PRJNA681031. Scripts used to calculate the amino acid compositional indices are available at https://github.com/agentilhomme/Genomic-Signatures-of-OGT-2020-.

## Author contributions

**Kari Green:** Formal analysis, Investigation, Visualization, Writing – original draft. **R. Eric Collins:** Conceptualization, Methodology, Software, Resources, Supervision, Project administration, Funding acquisition, Writing – review & editing.

## Supplementary Information

**Table S1:** CheckM completeness and contamination estimates for the MG-I genomes and MAGs analyzed in this study (BV-BRC genome IDs).

| Genome ID | Completeness (%) | Contamination (%) |
|---|---|---|
| 139207.4 | 99.5 | 0.5 |
| 228908.8 | 73.13 | 0 |
| 436308.17 | 100 | 0.97 |
| 517606.13 | 80.74 | 0.97 |
| 517606.5 | 95.63 | 7.61 |
| 660518.15 | 99.57 | 1.7 |
| 718286.4 | 100 | 0 |
| 859192.6 | 99.03 | 5.83 |
| 859350.6 | 92.39 | 1.94 |
| 886738.12 | 98.06 | 0 |
| 1001994.6 | 100 | 0 |
| 1229908.8 | 94.66 | 0 |
| 1229909.8 | 97.09 | 0 |
| 1379390.5 | 73.83 | 2.8 |
| 1470066.3 | 98.54 | 0 |
| 1470067.3 | 100 | 0 |
| 1580092.5 | 100 | 0 |
| 1582439.9 | 100 | 0.97 |
| 1655561.3 | 99.51 | 0.97 |
| 1671876.3 | 100 | 0 |
| 1776294.3 | 100 | 0 |
| 1898749.6 | 100 | 0 |
| 1915596.3 | 95.15 | 0 |
| 1915597.3 | 93.2 | 0 |
| 1915598.3 | 79.21 | 0 |
| 1915705.3 | 81.91 | 0 |
| 1959104.3 | 99.51 | 0 |
| 2022694.3 | 62.81 | 0.97 |
| 2022695.3 | 84.63 | 0.56 |
| 2022696.3 | 80.1 | 3.4 |
| 2022697.3 | 93.95 | 1.94 |
| 2024843.101 | 99.51 | 0.49 |
| 2024843.16 | 68.61 | 0.97 |
| 2024843.17 | 91.35 | 0.97 |
| 2024843.18 | 89.32 | 0 |
| 2024843.19 | 97.09 | 1.94 |
| 2024843.21 | 94.25 | 0 |
| 2024843.27 | 83.1 | 5.83 |
| 2024843.29 | 77.47 | 4.85 |
| 2024843.33 | 58.86 | 10.36 |
| 2024843.36 | 89 | 5.74 |
| 2024843.37 | 99.51 | 0.49 |
| 2024843.39 | 78.4 | 0 |
| 2024843.46 | 97.57 | 0 |
| 2024843.47 | 98.54 | 0.97 |
| 2024843.51 | 81.18 | 3.88 |
| 2024843.52 | 93.69 | 0.97 |
| 2024843.53 | 98.06 | 0 |
| 2024843.59 | 52.43 | 0 |
| 2024843.6 | 100 | 0 |
| 2024843.61 | 87.38 | 0.97 |
| 2024843.62 | 95.15 | 1.46 |
| 2024843.63 | 95.15 | 0.97 |
| 2024843.67 | 85.41 | 6.8 |
| 2024843.7 | 95 | 1.94 |
| 2024843.79 | 82.78 | 6.37 |
| 2024843.8 | 92.72 | 2.91 |
| 2024843.83 | 87.34 | 2.91 |
| 2024843.85 | 77.62 | 5.83 |
| 2024843.9 | 91.26 | 0.97 |
| 2024843.91 | 95.15 | 0 |
| 2024843.92 | 100 | 0.97 |
| 2024843.93 | 95.15 | 0 |
| 2024843.94 | 99.03 | 0 |
| 2026714.3 | 92.99 | 3.31 |
| 2026768.13 | 99.51 | 0.97 |
| 2026768.14 | 85.44 | 2.17 |
| 2026768.16 | 86.23 | 5.42 |
| 2026768.19 | 76.24 | 6.31 |
| 2026768.21 | 93.69 | 4.05 |
| 2026768.22 | 98.54 | 0.97 |
| 2026768.24 | 99.51 | 2.91 |
| 2026768.3 | 58.82 | 31.96 |
| 2026768.8 | 94.17 | 0 |
| 2026886.16 | 84.63 | 3.64 |
| 2026886.19 | 97.73 | 0.97 |
| 2026886.26 | 80.42 | 1.05 |
| 2026886.27 | 74.92 | 0.47 |
| 2026886.28 | 76.54 | 5.9 |
| 2026886.29 | 93.2 | 7.77 |
| 2026886.3 | 99.03 | 1.94 |
| 2026886.31 | 97.09 | 0 |
| 2026886.33 | 97.09 | 0 |
| 2026886.35 | 97.09 | 0 |
| 2026886.37 | 91.42 | 1.05 |
| 2026886.38 | 99.03 | 1.46 |
| 2026886.42 | 96.12 | 1.94 |
| 2026886.43 | 99.03 | 1.46 |
| 2026886.6 | 98.54 | 1.94 |
| 2026886.7 | 93.13 | 6.63 |
| 2041147.3 | 96.88 | 1.01 |
| 2109907.3 | 98.06 | 5.96 |
| 2109908.3 | 99.51 | 0.97 |
| 2109909.3 | 100 | 0 |
| 2202732.11 | 95.39 | 0 |
| 2202732.12 | 93.2 | 0 |
| 2202732.14 | 85.44 | 1.94 |
| 2202732.18 | 77.02 | 0.07 |
| 2202732.19 | 87.54 | 1.94 |
| 2202732.21 | 98.71 | 1.94 |
| 2202732.22 | 96.6 | 1.94 |
| 2202732.23 | 94.9 | 1.94 |
| 2202732.24 | 98.06 | 1.94 |
| 2202732.26 | 96.12 | 1.94 |
| 2202732.27 | 97.09 | 1.94 |
| 2202732.28 | 96.12 | 1.94 |
| 2202732.3 | 63.11 | 5.83 |
| 2202732.34 | 99.03 | 1.61 |
| 2202732.35 | 89.08 | 0.65 |
| 2202732.36 | 85.6 | 0 |
| 2202732.37 | 64.89 | 5.34 |
| 2202732.4 | 89.48 | 0 |
| 2202732.41 | 89.32 | 4.85 |
| 2202732.43 | 91.75 | 4.37 |
| 2202732.44 | 95.63 | 0.97 |
| 2202732.45 | 91.64 | 1.94 |
| 2202732.46 | 98.54 | 0.97 |
| 2202732.5 | 99.03 | 0.97 |
| 2202732.57 | 100 | 0.97 |
| 2202732.58 | 90.78 | 0 |
| 2202732.59 | 92.23 | 0.97 |
| 2202732.61 | 85.44 | 0.97 |
| 2202732.62 | 95.15 | 0.97 |
| 2202732.7 | 83.01 | 0.81 |
| 2202732.9 | 98.06 | 3.88 |
| 2249327.3 | 87.7 | 14.56 |
| 2259673.3 | 100 | 0 |
| 2478553.3 | 91.75 | 0.97 |
| 2478554.3 | 92.23 | 0 |
| 2478555.3 | 89.16 | 0 |
| 2478556.3 | 89.81 | 0.07 |
| 2478979.3 | 93.2 | 0 |
| 2508726.3 | 100 | 0.97 |
| 2511933.3 | 100 | 0 |
| 2795383.3 | 99.03 | 0 |

**Figure S1:**
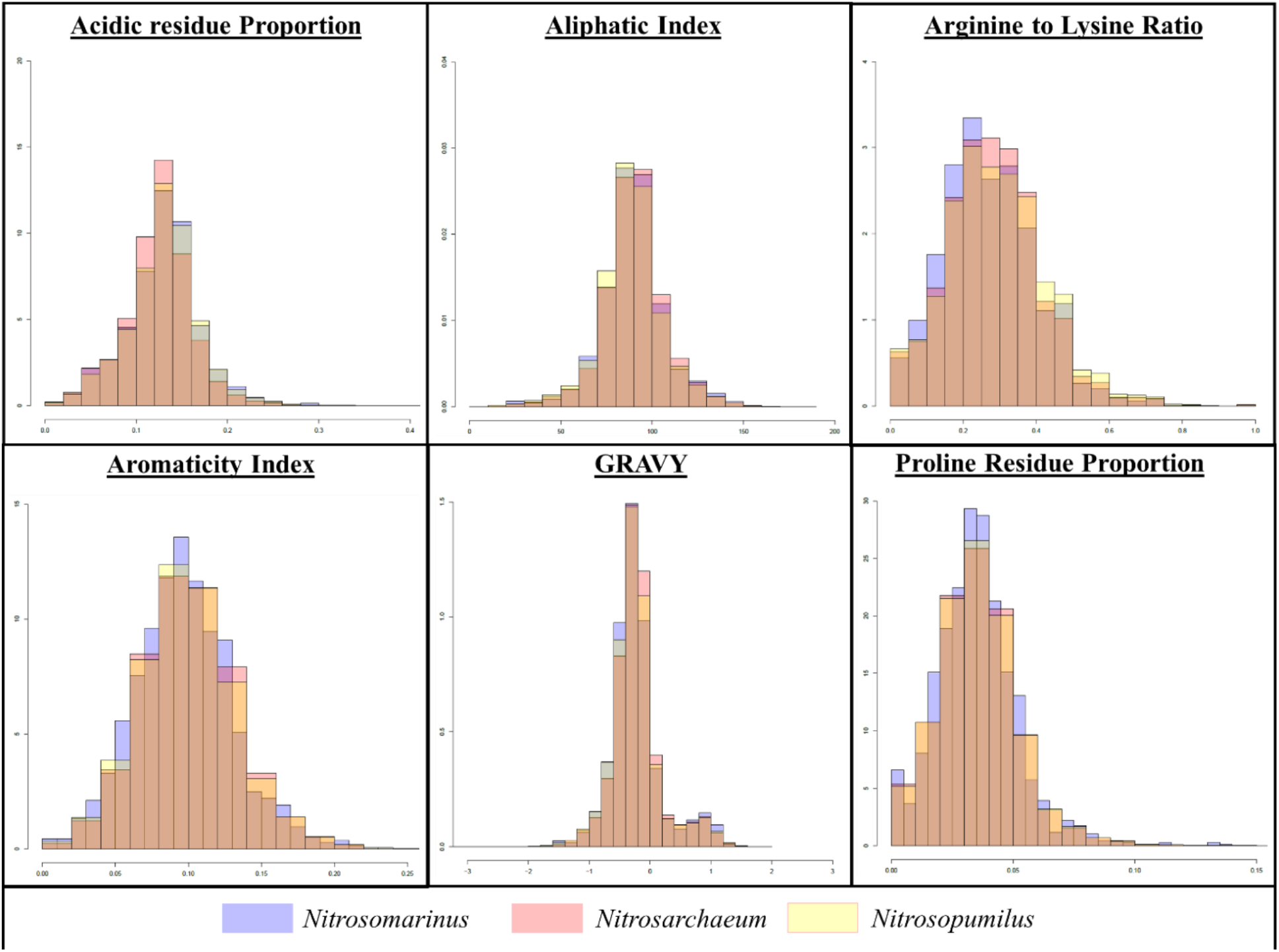
Amino acid indices distribution for the three MG-I genera analyzed.

**Figure S2:**
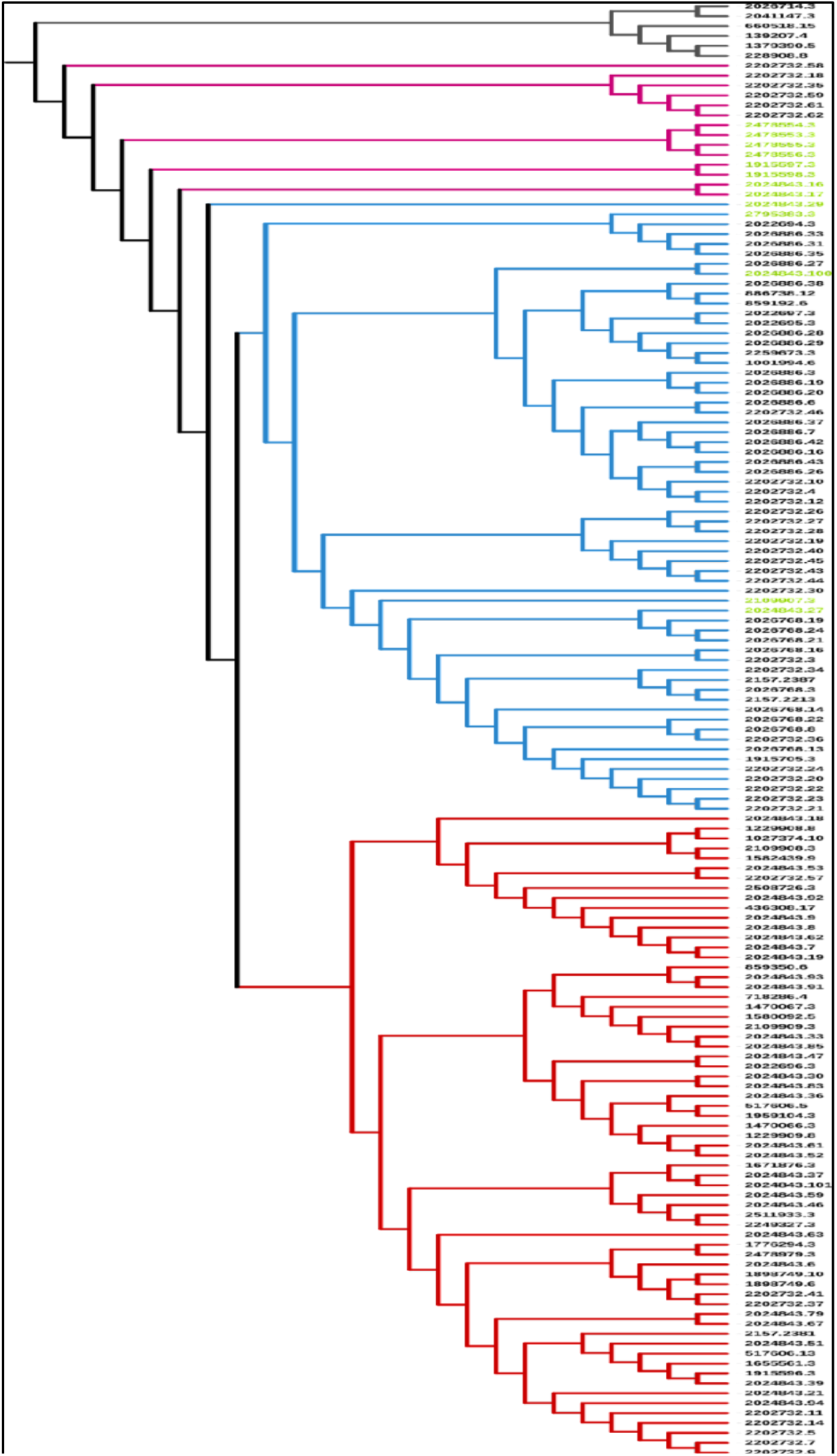
Annotated phylogenetic tree of the MG-I MAGs analyzed in Sections 3.1 and 3.3, including group A branches (red), group B branches (blue), sponge-associated genomes (green), Nitrosopumilaceae outliers (pink), and the outgroup (grey).

**Figure S3:**
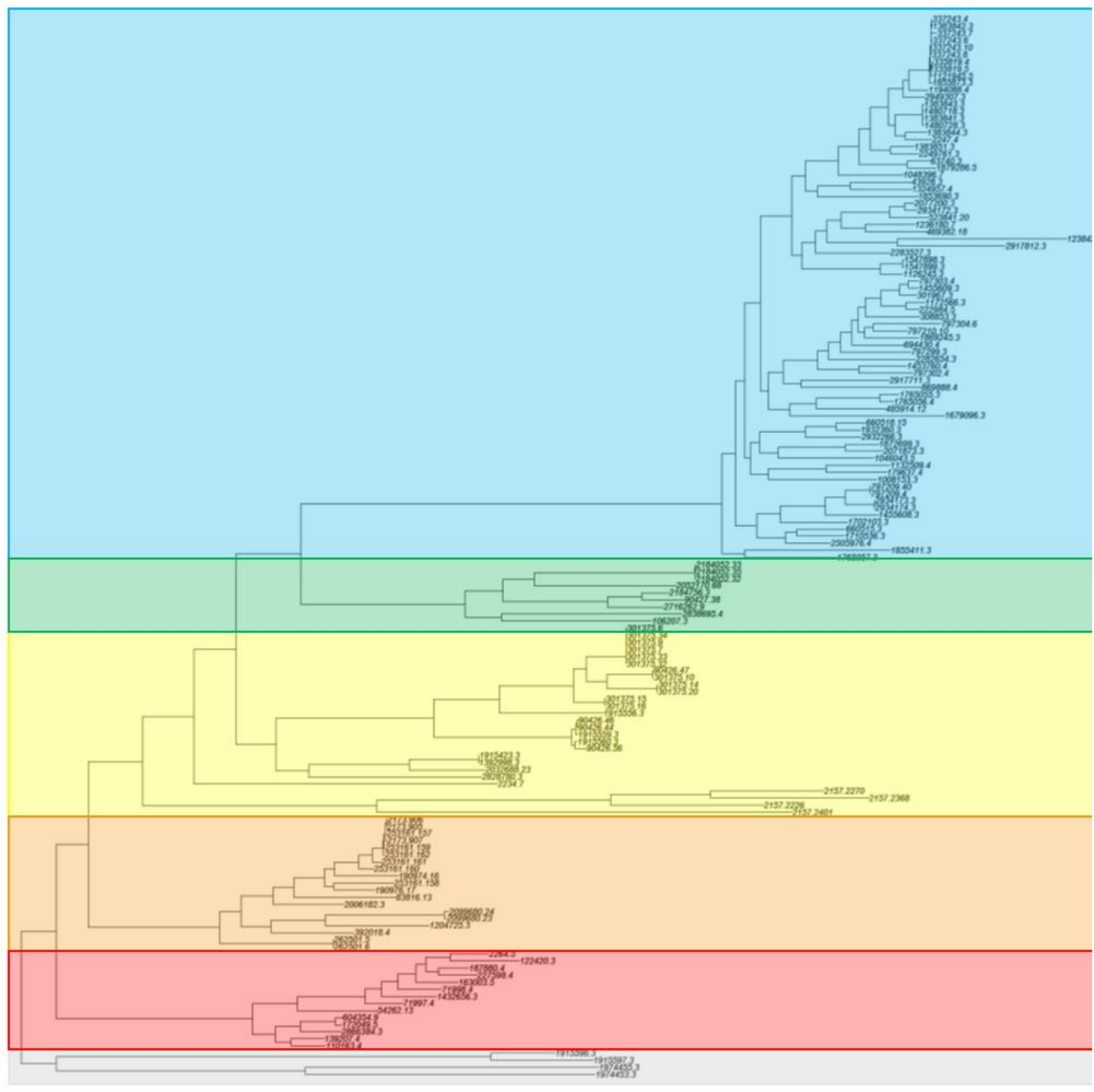
Annotated phylogenetic tree of the Euryarchaeota MAGs analyzed in Section 3.2, including Groups A (blue), B (green), C (yellow), D (orange), and E (red), and the outgroups (grey).

**Figure S4:**
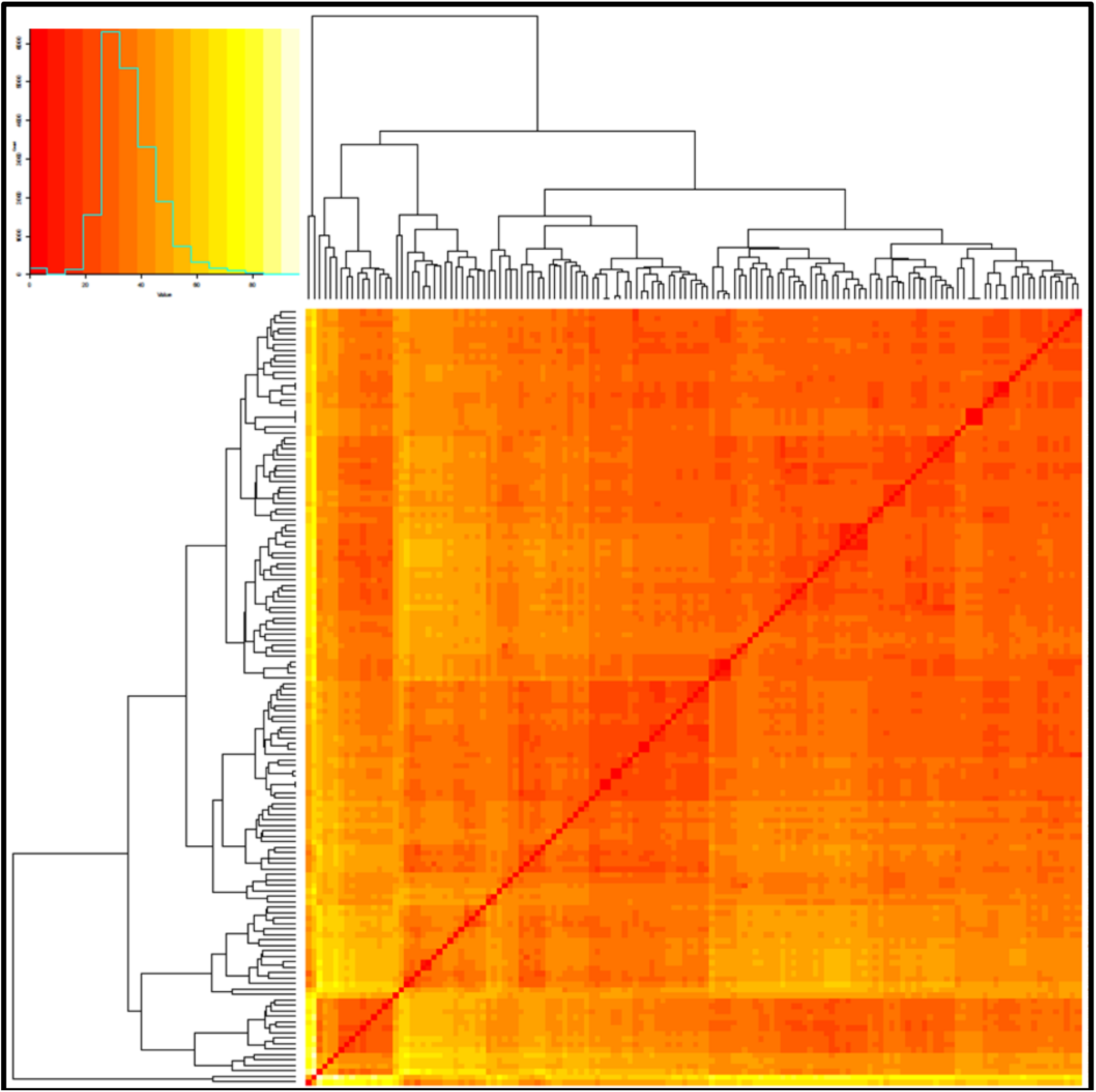
Pair-wise distances of the presence or absence of archaeal global protein families in the MG-I MAGs. Both y- and x-axes are the same, and the dark red line indicates a direct MAG to MAG comparison.

